# Algae-specific Immune Modulation Influences Responses to Heat and Pathogen Challenge in a Symbiotic Coral

**DOI:** 10.1101/2025.11.05.684850

**Authors:** Jeric Da-Anoy, Mu-Han Chen, Andrew Bouchie, John Dougherty, Anna K.H. Lapadula, Abigail Skena, Wei Wang, Timinte Abraham, Kian R. Thompson, Oliwia Jasnos, Kyle S. Toyama, Joshua Ayivor, Oyejadesola Diya, Thomas D. Gilmore, Sarah W. Davies

## Abstract

The role of symbiotic algae in coral life history and host health is well documented, but the immune and physiological trade-offs of hosting these symbionts remain less explored. While association with the algal symbionts of the genus *Durusdinium* is known to confer thermotolerance, it has also been linked to coral tissue loss under stress. We investigated whether algal type influences host immunity and stress responses in the tropical coral *Pocillopora acuta*. *Durusdinium*-hosting (*D*-hosting) *P. acuta* have distinct transcriptomic profiles, higher immune-related gene expression, and elevated baseline levels of immunity transcription factor NF-κB as compared to corals hosting *Cladocopium* (*C*-hosting). Under heat challenge, *D*-hosting *P. acuta* exhibited tissue loss, oxidative stress, immune and microbial dysregulation, whereas *C*-hosting *P. acuta* were more susceptible to bleaching, metabolic dysregulation, and decline in nitrogen-fixing and antioxidant-producing bacteria. Finally, infection with the the bacterium *Vibrio coralliilyticus* caused high tissue loss in *D*-hosting corals, but not in *C*-hosting corals. Our results suggest a mechanism for how *Durusdinium* association enhances thermotolerance yet predisposes corals to tissue damage under stress, suggesting immune trade-offs that can compromise host survival under multiple stressors.

## INTRODUCTION

Symbiosis between coral hosts and obligate intracellular algae in the family Symbiodiniaceae (*1*) enables them to flourish in tropical reef waters (*2*). In a stable symbiosis, algae can translocate up to 95% of their photosynthetically fixed carbon to the host, supplying the host’s metabolic needs (*3*). However, under elevated temperatures, algal photosystems can be disrupted (*4*) and trigger an overproduction of reactive oxygen species (ROS) (*5*), which can overwhelm host antioxidant defenses (*6*), induce lipid peroxidation of cell membranes (*7*), and eventually induce symbiont expulsion in a process called coral bleaching or dysbiosis (*8*). This breakdown in symbiosis can be influenced by host genetic background (*9*), algal associations (*10*), microbiome communities (*11*), and environmental history (*12*), making predicting coral bleaching a challenge.

One molecular process involved in the maintenance of cnidarian-algal symbiosisis the downregulation of host immune pathways (*13*), which could have implications for host susceptibility to secondary biotic or abiotic stresses. For example, in the sea anemone *Exaiptasia diaphana* (Aiptasia), symbiotic anemones infected with either *Pseudomonas aeruginosa* or *Serratia marcescens* exhibit lower survival rates as compared to their aposymbiotic counterparts (*14*). This pattern has also been observed in polyps of the upside-down jellyfish *Cassiopea xamachana* infected with *Serratia marcescens* (*15*). However, specific algal associations can also impact other host responses (*16*). For example, *Pocillopora damicornis* associated with algal symbiontis of genus *Durusdinium* (formerly Clade D) are more resistant to thermal challenge than those hosting *Cladocopium* (formerly Clade C) (*17*). This same pattern of coral-algae associations has been observed in natural marine settings in response to heatwaves. Before a heatwave in the Persian/Arabian Gulf, *Pocillopora spp.* were most often associated with *Cladocopium,* but after the heatwave most surviving corals were hosting *Durusdinium* (*18*), suggesting that *Durusdinium* conferred a survival advantage under conditions of elevated water temperatures. However, there may be negative impacts to hosting *Durusdinium.* For example, although *D*-hosting *Montipora capitata* were more resistant to bleaching than *C*-hosting conspecifics on the same reef, *D*-hosting conspecifics were more susceptible to disease or tissue loss (*19*). Taken together, such results suggest that while hosting *Durusdinium* can enhance thermotolerance, its association can compromise host health and defense to a secondary threat, such as pathogen infection.

While host immunity and symbiont associations have both been shown to influence coral health and stress tolerance, the molecular basis of their interactions is not fully understood. For example, how symbiont type affects immune regulation is not fully described. Transcription factor NF-κB is a key regulator of immune responses to microbial infections and environmental stress in many organisms (*20*). Previous studies have demonstrated that baseline immune status, including NF-κB pathway genes, varies between symbiotic states in two facultative cnidarians (*13*, *21*) and that heat stress induces an innate immune response during coral bleaching (*22*), which is often associated with upregulation of NF-κB (*23*). Given that suppression of NF-κB is linked to the establishment and maintenance of symbiosis (*13*), the levels of NF-κB in a cnidarian host may depend on the type of algal associations it is undergoing, potentially linking stress susceptibility to differences in NF-κB levels between *C*- and *D*-hosting corals. Indeed, in the sea anemone Aiptasia the expression of NF-κB protein is suppressed in the presence of the native symbiont, *Breviolum* strain SSB01, but not by several *Durusdinium* strains (A001, Mf2.2b, and Ap2) (*24*). However, the mechanistic interactions between symbiont-mediated modulation of NF-κB and its consequences for holobiont stress responses remain largely unexplored, especially in tropical stony corals.

Here, we used *Pocillopora acuta* colonies associated with either *Cladocopium*, an algal symbiont that has co-evolved with other Pocilloporid corals *(P. meandrina, P. grandis, P. verrucosa)* (*25*), or *Durusdinium*, an algal symbiont that is generally associated with *Pocillopora spp.* exposed to high water temperatures or turbid environments (*26*), to address two questions: 1) Do these distinct algal symbiont associations impact baseline host immune status, and if so, 2) How do these immunological differences impact coral host responses to stress? Through the integration of physiological, cellular, and molecular approaches, our results reveal how algal symbiont-mediated immune differences shape coral stress tolerance, uncovering key pathways associated with coral responses to warming waters.

## RESULTS

### *Durusdinium* association is linked to increases in both immune gene expression and NF-κB levels in *Pocillopora acuta*

To characterize baseline gene expression differences between *C*- and *D*-hosting *P. acuta*, we conducted TagSeq gene expression profiling on three replicates of five host:symbiont pairings (1:C, 2:D, 3:D, 4:C, 5:D). PCA of all host genes revealed significant clustering between *C*- and *D*-hosting corals along PC1, which explained 52.3% of the variation in expression (P_Symbiont_<0.001), and additional clustering by host genotype (P_Genotype_<0.001) (Fig. 1A). Differential gene expression analysis identified 442 upregulated and 375 downregulated genes in *D*-hosting as compared to *C*-hostin*g* corals (Fig. 1B, P_Adj_ < 0.05). Gene Ontology (GO) enrichment analysis showed that *Biological Process* terms related to immunity were enriched in *D*-hosting compared to *C*-hosting corals, which included immune response [GO:0006955], immune effector process [GO:0002252], and cell activation involved in immune response [GO:0002263] (Fig. 1C, Mann-Whitney U (MWU) test p<0.05). When Differentially Expressed Genes (DEG) within these GO terms were explored, *C*-hosting corals showed reduced expression of classic immune-related genes relative to *D-*hosting corals (e.g., *Catalase/Cat*, *Peroxidasin/PXDN*, *Tumor Necrosis Factor Receptor-Associated Factor/TRAF*, *Interferon Regulatory Factor*/*IRF*, p_adj_<0.05) (Fig. 1D). Western blotting of four replicates of three host:symbiont pairings revealed that *D*-hosting corals exhibited 4-fold (2:D) and 4.1-fold (3:D) higher levels of full-length Pa-NF-κB transcription factor compared to the *C*-hosting coral (P_Genotype_<0.001, Fig. 1D), and these differences were not due to to differences in total protein loading (as judged by Ponceau staining). As a control for the cross-reactivity of the antiserum, we show that the anti-Aiptasia NF-κB antiserum used here can specifically detect a protein of the correct size in *P. acuta* extracts (fig. S1). That is, an approximately 100 kDa band detected in *P. acuta* extracts co-migrates with the primary band detected in extracts from HEK 293T cells transfected with an expression plasmid for *P. acuta* NF-κB (Pa-NF-κB).

**Fig. 1.**
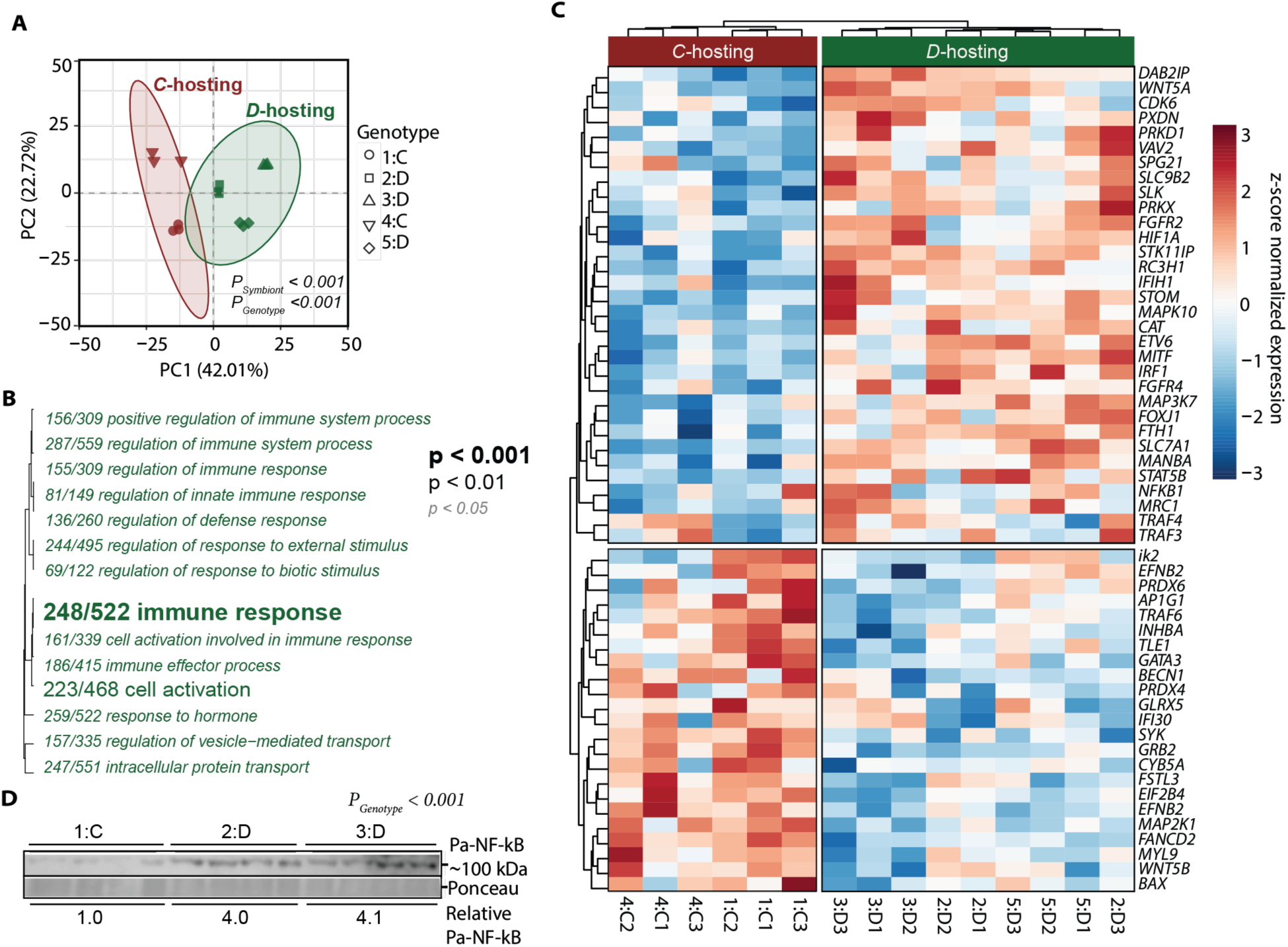
Distinct immune profiles associated with *C*- and *D*-hosting corals. (**A**) PCA of all host genes showing distinct clustering by algal type. Ellipses denote the 95% confidence intervals and p-values for algal type and host genotype are shown with axis percentages representing the total explained variance. (**B**) Gene ontology (GO) *Biological Process* terms enriched in *D-*hosting (green) versus *C*-hosting corals. Font size depicts the significance level as indicated by the inset key, and the proportion preceding each GO term designates the number of genes passing a raw p-value <0.05 out of the total genes belonging to that GO term. Linkage between terms, indicated by tree, shows relatedness of GO terms based on shared genes. (**C**) Heatmap showing expression of genes belonging to immunity-related GO terms across *C*- and *D*-hosting corals. Each row is a gene and columns are samples. Color of the cell represents row-scaled expression values (z-scores), indicating the relative difference in expression for each gene from its mean across all samples (blue, downregulated; red, upregulated). (**D**) Western blot and quantification of Pa-NF-κB levels in *D*-hosting compared to *C*-hosting corals. To obtain the relative NF-κB levels, Pa-NF-κB levels in each lane were obtained by first quantifying the ratio of bands for Pa-NF-κB/Ponceau for each lane, and the three values for *C*-hosting or *D*-hosting lanes were then averaged. The average of NF-κB in the *C*-hosting lanes was normalized to 1.0 and the average of the *D*-hosting corals is relative to the value for the *C*-hosting lanes. P_Genotype_ indicates significant differences across host:symbiont pairings (*p<0.001*).

### Conservation of activities in the *P. acuta* transcription factor NF-κB

Because NF-κB protein levels were different in *C*- vs. *D*-hosting *P. acuta*, we next sought to determine the properties of the *P. acuta* NF-κB protein. Manual searching of genomic and transcriptomic databases identified a single full-length Pa-NF-κB protein of 919 amino acids (aa) with an N-terminal Rel Homology DNA-binding-dimerization domain (RHD), followed by a Gly-rich region (GRR), and a C-terminal domain that contained six ankyrin (ANK) repeats and a set of three serines for possible phosphorylation by IκB kinase (Fig. 2A and fig. S2, A and B). Thus, the domain structure of Pa-NF-κB is essentially identical to human NF-κB p100 as well as to Aiptasia NF-κB (*13*), Moreover, an AlphaFold3-generated model of the RHD sequences of Pa-NF-κB on a canonical DNA site (5’GGGAATTCCC3”) produced a homodimer with a structure that is similar to models of mouse NF-κB p50 and *Nematostella vectensis* NF-κB dimers on the same site (fig. S3A).

**Fig. 2.**
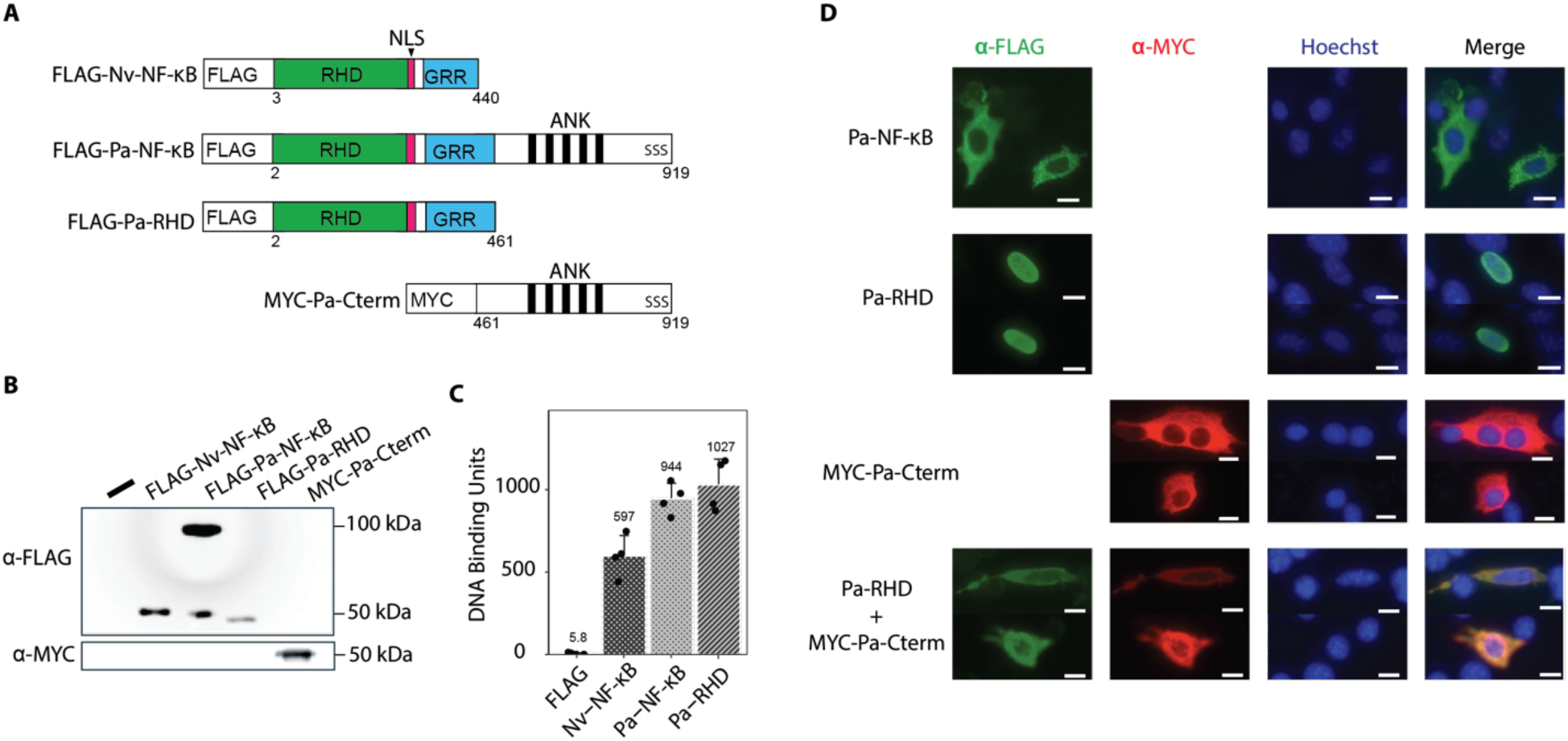
Characterization of the *P. acuta* transcription factor NF-κB. (**A**) General structures of NF-κB proteins, with N-terminal protein tags, used in these studies. RHD, Rel homology domain; NLS, nuclear localization sequence; GRR, glycine-rich region; ANK, ankyrin repeats (black bars). (**B**) Anti-FLAG and anti-MYC Western blotting of extracts from 293T cells transfected with expression plasmids for the indicated proteins. -, empty vector control. Molecular weight markers are indicated to the right. (**C**) DNA-binding units from 293T cell extracts (from **B**) incubated with an NF-κB binding site and analyzed by ELISA (see Materials and Methods). DNA-binding values represent averages of four samples. (**D**) Indirect immunofluorescence images from chicken DF-1 fibroblasts transfected with expression plasmids for indicated proteins and then stained for the indicated protein tags or Hoechst nuclear dye. White scale bar is 10 µm.

To analyze the activities of the Pa-NF-κB protein, we created FLAG-tagged expression vectors for full-length Pa-NF-κB (aa 2-919), an RHD-GRR (aa 2-461) protein, and a MYC-tagged version of the C-terminal ANK repeat domain (aa 461-919). As a control, we also used an expression vector for FLAG-Nv-NF-κB, the Nematostella NF-κB protein that we have characterized previously (*27*). These plasmids were transfected into HEK 293T cells, and anti-FLAG and anti-MYC Western blotting of extracts from these cells showed that the plasmids directed the expression of proteins of the appropriate sizes (Fig. 2B). Of note, there is also a minor amount of an anti-FLAG-reactive ∼50 kDa protein following transfection with full-length FLAG-Pa-NF-κB, which is presumably a C-terminally processed form of FLAG-Pa-NF-κB.

To analyze the DNA-binding activity of the FLAG-tagged proteins, we used whole-cell extracts from the transfected 293T cells in an ELISA-based κB-site DNA-binding assay that we recently developed (*28*). In these assays, extracts expressing full-length Pa-NF-κB and Pa-RHD-GRR contained ∼175-fold higher levels of κB-site DNA-binding activity, as compared to extracts from cells transfected with the vector control (Fig. 2C). DNA-binding activity levels in the extracts expressing Pa-NF-κB and Pa-RHD-GRR proteins were similar to the levels seen in extracts from cells expressing Nv-NF-κB (440 aa, positive control). These results indicate that the Pa-NF-κB protein can bind to a consensus NF-κB DNA site in vitro, even in the presence of the C-terminal ANK repeat sequences.

To characterize the subcellular localization of the full-length Pa-NF-κB, Pa-RHD-GRR, and Pa-Cterm proteins, we transfected the expression plasmids into chicken DF-1 fibroblasts and performed anti-FLAG or anti-MYC (for Pa-Cterm) indirect immunofluorescence (Fig. 2D). We found that full-length Pa-NF-κB and the Pa-Cterm protein were both localized exclusively in the cytoplasm. Conversely, the Pa-RHD-GRR protein was localized exclusively in the nuclei of cells. These results suggest that the C-terminal ANK repeat sequences can block the nuclear localization of the Pa-NF-κB protein. Moreover, when we co-transfected chicken DF-1 cells with the separate Pa-RHD-GRR and Pa-Cterm expression vectors, both proteins were now localized in the cytoplasm (Fig. 2D), suggesting that the Pa-Cterm sequences were interacting with the Pa-RHD-GRR protein in trans to block its nuclear localization.

Next, we investigated whether the C-terminal sequences of Pa-NF-κB can biochemically interact with the RHD sequences. We co-transfected the FLAG-tagged Pa-RHD-GRR plasmid and the MYC-tagged Pa-Cterm into 293T cells, lysed the cells, and then performed anti-FLAG co-immunoprecipitations. In samples from 293T cells co-transfected with the FLAG-tagged Pa-RHD-GRR plasmid and the MYC-tagged Pa-Cterm, both FLAG-Pa-RHD-GRR and MYC-Pa-Cterm were present in anti-FLAG immunoprecipitates. In contrast, MYC-Pa-Cterm was not pulled down in an anti-FLAG co-immunoprecipitation if it was co-transfected with the empty FLAG vector (fig. S3B). This result suggests that the Pa-RHD and C-terminal ANK repeat sequences of Pa-NF-κB can directly interact with one another in cells.

We further noted (fig. S2A) that Pa-NF-κB has a set of three serine residues (D<u>S</u>GLF<u>S</u>Q<u>S</u>) that are similar to ones that can lead to C-terminal processing of human and cnidarian NF-κB proteins (*29*). Namely, we have previously shown that co-expression of an IκB kinase (IKK) protein can induce the processing of both a coral and sea anemone NF-κB protein in human 293T cells (*13*, *29*). To determine whether IKK expression could induce processing of full-length Pa-NF-κB, we co-transfected human 293T cells with expression vectors for FLAG-Pa-NF-κB and a constitutively active human IKKβ protein (IKKβ-SS/EE). Co-expression of IKKβ-SS/EE led to a 2.6-fold increase in the amount of processed (∼50 kDa) Pa-NF-κB, as compared to cells co-transfected with Pa-NF-κB and the empty vector control (fig S3C). This result shows that co-expression of an active IKK protein can induce processing of Pa-NF-κB, at least in human cells.

Taken together, the results in this section indicate that the Pa-NF-κB protein has many of the structural, regulatory, and activity properties of mammalian and sea anemone NF-κB proteins.

### Algal symbiont identity results in different physiological responses to thermal challenge

To measure physiological responses of *C*- and *D*-hosting corals to thermal challenge, we conducted a 12-day common garden thermal challenge experiment on three host:symbiont pairings (1:C, 2:D, 3:D). Overall, the photophysiology of the *C*-hosting genotype was more negatively impacted by heat challenge than the two *D*-hosting genotypes. The magnitude of heat-induced photosynthetic efficiency (F_v_/F_m_) declines differed significantly among genotypes (*X*^2^ = 31.536, df = 2, p < 0.001), with 1:C showing greater declines over time compared to the two *D*-hosting genotypes (fig. S4, Table S1). Specifically, we observed divergence in responses on the final day of thermal challenge (Fig. 3B), with an approximately 3.4-fold greater reduction experienced by the *C*-hosting genotype (1:C: mean decrease: 0.439 ± 0.039, t(10) = 11.185, p < 0.001), compared to the two *D*-hosting genotypes (2:D: mean decrease: 0.130 ± 0.039, t(10) = 3.313, p = 0.008; 3:D: mean decrease: 0.249 ± 0.039, t(10) = 6.338, p < 0.001; Table S2). Red channel intensity, a proxy for bleaching, increased significantly under heat challenge for all corals (*X*^2^ = 58.066, df = 1, p < 0.001); however, the magnitude of bleaching differed among genotypes (*X*^2^ = 25.278, df = 2, p < 0.001) with 1:C showing the greatest change (mean increase: 105.0 ± 13.8%, t(11) = 7.620, p < 0.001) compared with the two *D-*hosting genotypes (2:D: mean increase: 31.2 ± 13.8%, t(11) = 2.265, p = 0.045; 3:D: mean increase: 28.6 ± 13.8%, t(11) = 2.073, p = 0.062; Fig. 3D, Table S3).

**Fig. 3.**
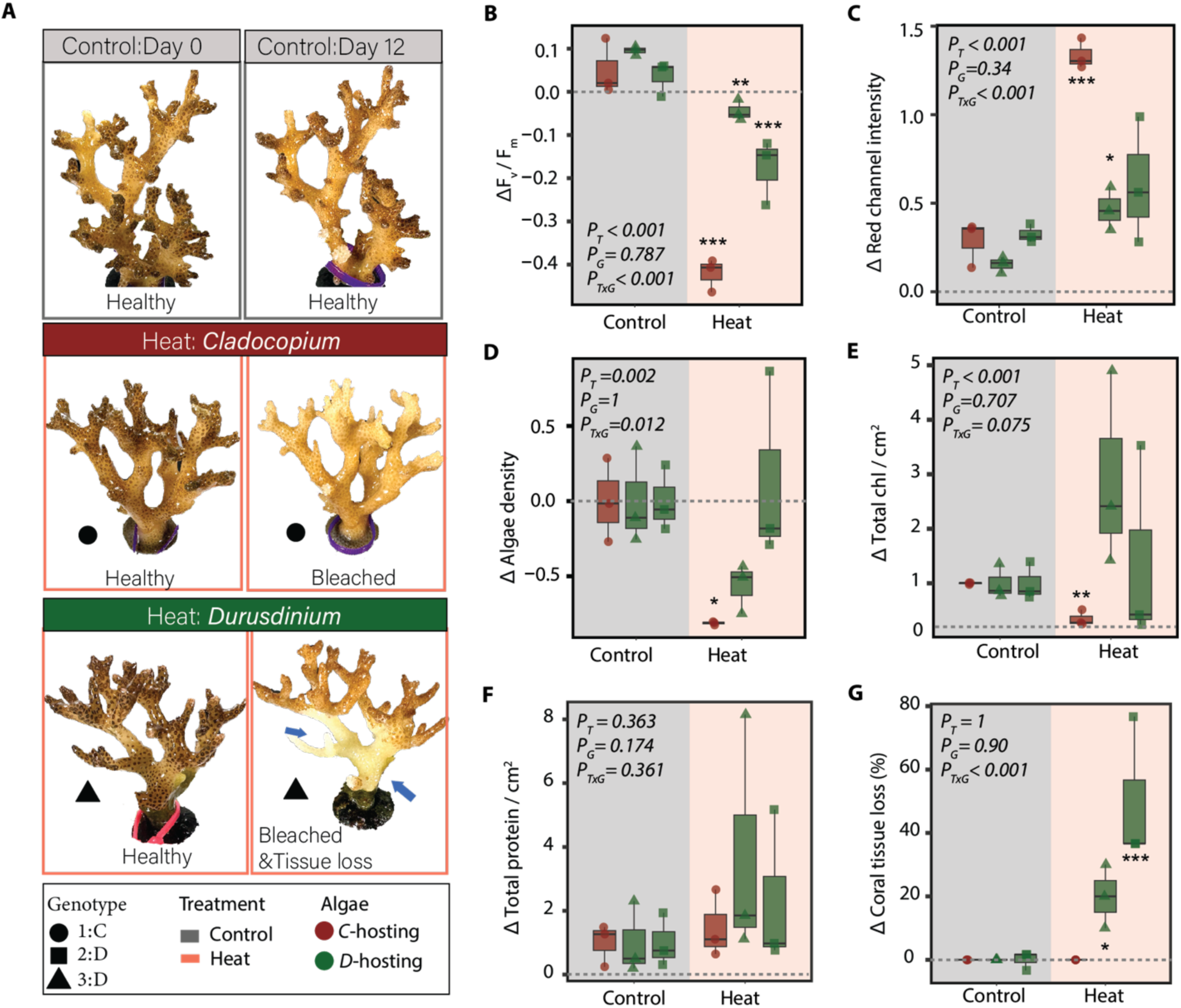
Physiological responses of *C-* and *D*-hosting corals in response to heat challenge. (**A**) Representative photographs of coral fragments before and after heat challenge. Fragments exhibited various phenotypes under heat challenge: healthy, bleached, and tissue loss (blue arrows).(B) Change in photosynthetic efficiency (ΔF_v_/F_m_), (C) change in red channel intensity calculated as (Day 12 - Day 0) / Day 0 (Δ red channel intensity), and (D) relative change in algal density (Δ algal density). (**E**) relative change in Total chlorophyll (chlorophyll *a* + *c*) and (**F)** relative change in total protein yield of *C-* and *D*-hosting corals after heat challenge at the end of the experiment. In (**B-F**) P_Treat_, P_Algae_, P_TxA_ indicate significance for heat challenge, algal type, and their interaction. Asterisks (***, **) indicate significant differences (p<0.001, p<0.01) between heat treatment and controls using Tukey’s HSD tests. (**G**) Percentage of tissue loss at day 12 under heat challenge. Tissue loss was never observed for *C*-hosting corals. Values in (**B**-**G**) are normalized relative to the control average for each genotype to account for differences in initial physiological status between *C*-hosting and *D*-hosting corals. In (**B-G**) Each point represents an individual coral fragment and shapes represent different host:symbiont pairings: 1:C (circle), 2:D (square), and 3:D (triangle), background color represents control (grey) and heat (pink) treatments, while box plot color represents algal symbiont type (brown = *C*-hosting, green = *D*-hosting).

Algal density was also significantly reduced under heat challenge for all corals (*X*^2^ = 8.963, df = 1, p = 0.003); however, declines were significantly greater in the *C*-hosting genotype (1:C: mean decrease: 81.2 ± 27.1%, t(10.6) = 2.994, p = 0.013) relative to the two *D-*hosting genotypes where cell densities decreased in 2:D under heat challenge (mean decrease: 56.4 ± 27.1%, t(10.6) = 2.077, p = 0.063), and no change was observed in 3:D (mean increase: 13.1 ± 27.1%, t(10.6) = −0.484, p = 0.639; Fig. 3C, Table S4). Heat challenge significantly reduced total chlorophyll in all corals (*χ*^2^ = 59.082, df = 1, p < 0.001), with responses differing marginally among genotypes (*X*^2^ = 5.179, df = 2, p = 0.075). Specifically, 1:C was the only pairing that showed a significant decrease under heat challenge (mean decrease: 1.12 ± 0.146 µg cm^−2^, t(4) = 7.686, p = 0.002), while 2:D (mean decrease: 5.92 ± 3.240 µg cm^−2^, t(4) = 1.823, p = 0.142) and 3:D (mean decrease: 1.26 ± 3.400 µg cm^−2^, t(4) = 0.370, p = 0.730) showed no significant changes (Fig. 3E, Table S5).

Protein content did not differ between treatments (*χ*^2^ = 0.032, df = 1, p = 0.857) or genotypes (*X*^2^ = 0.32, df = 2, p = 0.850; Fig. 3F, Table S6).

Tissue loss patterns differed significantly among genotypes under heat challenge (*X*^2^ = 19.913, df = 2, p < 0.001), with 1:C showing no tissue loss (mean difference: 0 ± 7.98%, t(10) = 0.000, p = 1.000), 2:D experiencing moderate tissue loss (mean difference: 20.0 ± 7.98%, t(10) = 2.508, p = 0.031), and 3:D showing the most extensive tissue loss (mean difference: 50.0 ± 7.98%, t(10) = 6.269, p < 0.001; Fig. 3G, Table S7–8).

### Weighted gene co-expression network analysis (WGCNA) reveals distinct heat-responsive processes between algal associations

To investigate molecular processes associated with the different physiological responses of *C-* and *D-*hosting corals, we profiled gene expression of *P. acuta* fragments from the three host:symbiont pairings (1:C, 2:D, 3:D) at the end of the heat challenge experiment, compared these patterns to baseline conditions, and then assessed functional responses using WGCNA and gene ontology (GO) enrichment analyses. The identified modules are linked to a variety of treatments and physiological traits (fig. S5); however, the cyan module (2049 genes, R²= 0.82 in heat, R²= 0.76 in sloughing, R²= 0.58 in red channel intensity) showed the highest positive correlation with heat treatment. GO enrichment of *Biological Processes* in the cyan module revealed significant overrepresentation of signaling pathways known to maintain cell integrity and symbiosis under heat challenge. This enrichment included immune response [GO:0002376, GO:0002699, GO:0002218, GO:0002758], apoptosis [GO:0032680, GO:0034612, GO:0071356], and regulation of the mitogen-activated protein kinases (MAPK) cascade [GO:0043408, GO:0032874, GO:0070304, GO:0046330] (Fig. 4A).

**Fig. 4.**
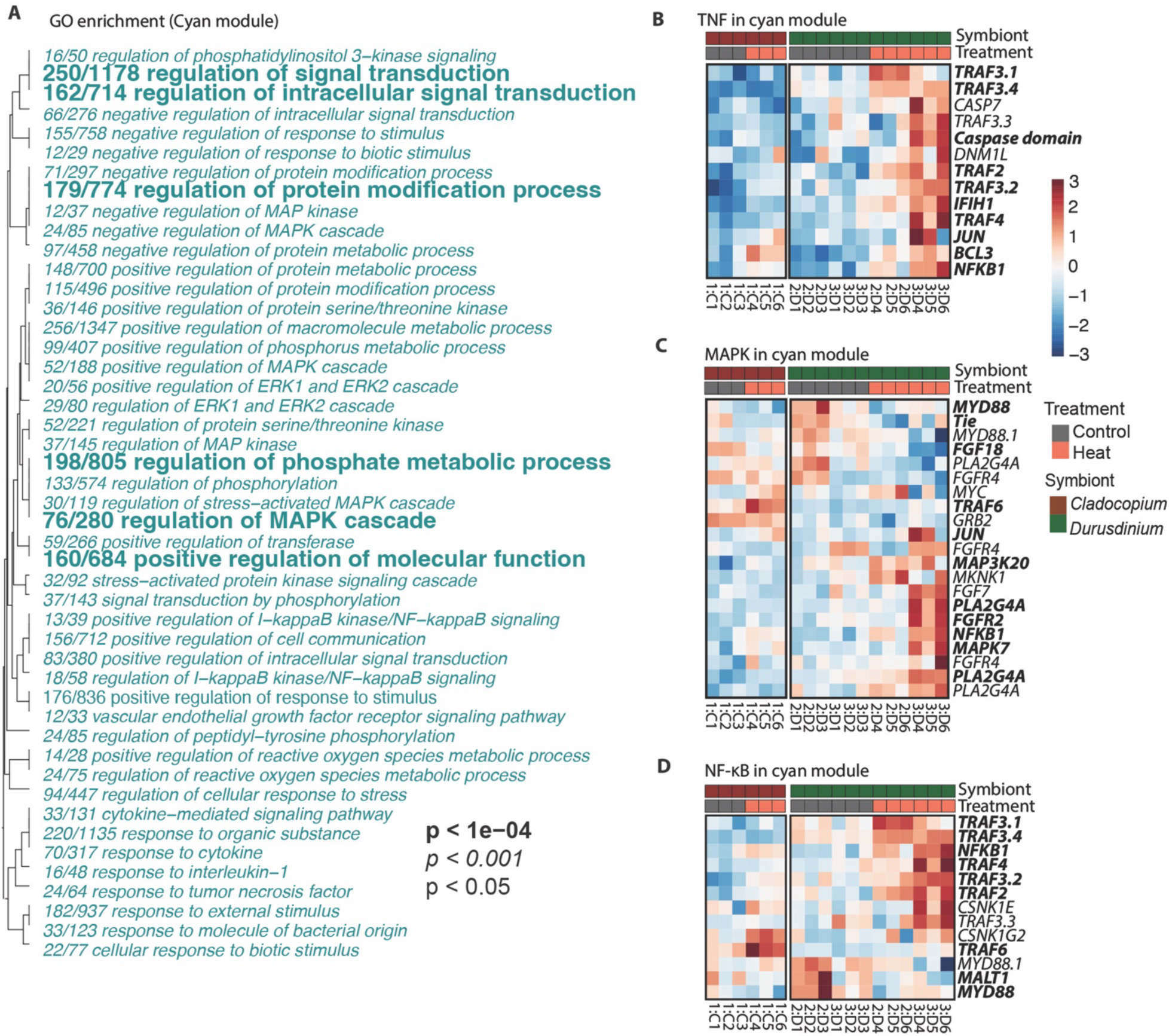
Weighted Gene Co-expression Network Analysis (WGCNA) reveals immune and stress response pathways associated with heat treatment and physiological traits. (**A**) Gene Ontology enrichment of genes assigned to the cyan module (N=2,049 genes), which showed positive correlations with heat treatment, tissue sloughing, and color loss (see fig. S5 for WGCNA results). (**B-D**) Heatmaps of genes enriched in the cyan module associated with (**B**) TNF signaling, (B) MAPK, and (**D**) NF-κB. Bold gene names denote genes that were significantly differentially expressed (p_adj_ < 0.05). The gene encoding NF-κB(*NFKB1*) appears in **B-D** given its role as a downstream effector of both TNF and MAPK signaling. Each row is a gene and columns are samples. Cell color represents row-scaled expression values (z-scores), indicating the relative difference in expression for each gene from its mean across all samples (blue, downregulated; red, upregulated).

We then examined some immune GO terms known to be associated with maintaining and regulating symbiosis (*30*). DEGs involved in Tumor Necrosis Factor (TNF), MAPK, and NF-κB pathways were manually searched for in the cyan module (Fig. 4B, C, and D). Heat challenge resulted in increased expression of the major players of the MAPK pathway (*MAPK7*, *FGF4*, and *MAP3K20*) and TNF (*TRAF3*, *IFIH1*, *CASP7*, *NFKB1*, *BCL3*, and *DNM1L*) pathways and under heat challenge these genes were more highly expressed in *D*-hosting than *C-*hosting corals (Fig. 4B and C, Table S9, 10, and 11). Notably, genes associated with the NF-κB pathway (*NFKB1*, *TRAF3*) exhibited higher expression under heat challenge, and expression was particularly high in *D-*hosting corals (Fig. 4D, Table S11).

### *D*-hosting corals show greater upregulation of stress- and symbiosis-related genes under thermal challenge

To determine whether common heat stress genes and symbiosis-associated genes are differentially regulated under heat in *D*-hosting (2:D, 3:D) and *C*-hosting corals (1:C), we analyzed gene expression patterns of known heat stress response and symbiosis genes. Comparison of heat-responsive genes in *C*-hosting and *D*-hosting corals revealed that cellular stress response pathways, including heat shock proteins (*HSPA5*, *HSP90B1*, *HSPA8*, *HSPA4L*), immune modulators (*NFKB1*, *BCL3*, *TRAF3.1*, *TRAF3.2, TRAF3.4, TRAF4*), and molecular mediators such as *MIB2* and *CYP21A2* were upregulated under heat treatments (Fig. 5A, table S12). Moreover, *D*-hosting corals exhibited higher induction of these genes, as well as antioxidant genes (*Peroxidases*, *Glutathione S-Transferase*), relative to the *C*-hosting coral (Fig. 5A, table S12), suggesting increased plasticity of these genes in *D-*hosting corals under heat treatment. Similarly, symbiosis-related genes also differed between treatments (Fig. 5B, table S5). Corals under heat treatment upregulated numerous genes involved in symbiotic interactions, such as *PLA2G4A*, *ALOX5*, *ALDH3B1*, *NPC2*, *swt*, and *SLC* transporters (Fig. 5B) and expression of symbiosis-related genes in the *C*-hosting coral was relatively stable across treatments. In contrast, *D*-hosting corals exhibited greater upregulation of symbiosis-related genes under heat treatment relative to the *C*-hosting coral (Fig. 5B, table S13). These results show that *D*-hosting corals exhibit increased overall gene expression inducibility of cellular stress responses and symbiosis-related genes under heat treatment as compared to the *C*-hosting coral.

**Fig. 5.**
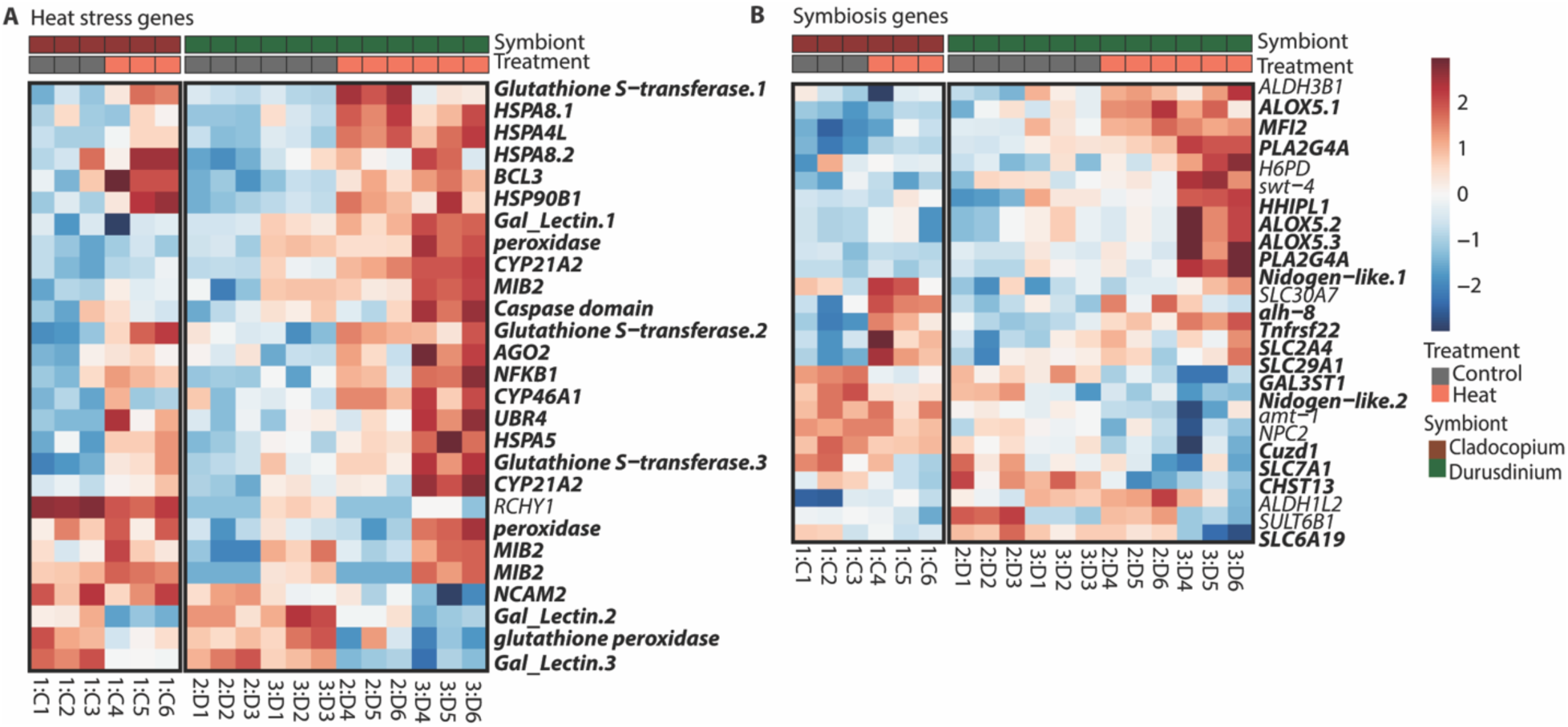
Heat stress and symbiosis-related gene expression vary under heat challenge and by algal type. (**A**) Heatmap of differentially expressed heat stress-related genes from (*31*) in *C*- (1:C) or *D-*hosting corals (2:D, 3:D) under control (grey) and heat challenge (orange). Genes shown include those involved in oxidative stress responses, immune signaling and protein regulation. (**B**) Heatmap of differentially expressed symbiosis-related genes from (*31*) under thermal challenge. Each row is a gene and columns are samples. Cell color represents row-scaled expression values (z-scores), indicating the relative difference in expression for each gene from its mean across all samples (blue, downregulated; red, upregulated).

### Minor shifts of coral microbiome communities across genotypes under thermal challenge

To explore the role of the coral microbiome in the physiological and molecular changes we observed between *C*-hosting and *D*-hosting corals under heat challenge, we used 16S rRNA gene sequencing of three host:symbiont pairings (1:C, 2:D, 3:D) exposed to increased temperature for eight days as compared to controls. Our results showed that heat challenge had a marginally significant effect on microbiome community structure (ADONIS F=1.95, R^2^=0.063, P=0.055, Fig. 6A, fig. S6A) and did not impact microbial alpha diversity (Shannon index F=0.09, P=0.77, evenness F=0.92, P=0.35; Fig. 6B; inverse Simpson F=1.31, P=0.27; ASV richness F=2.12, P=0.17; fig. S6B and C). Notably, microbiome communities were significantly structured by host genotype (ADONIS F=6.74, R^2^=0.439, p<0.001, Fig. 6B). These host genotype effects were also reflected in Shannon index (F=6.57, P=0.01) and evenness (F=11.35, P=0.002, Fig. 6B), suggesting host-specific differences in microbiome communities when they host *C* vs *D* algae.

**Fig. 6.**
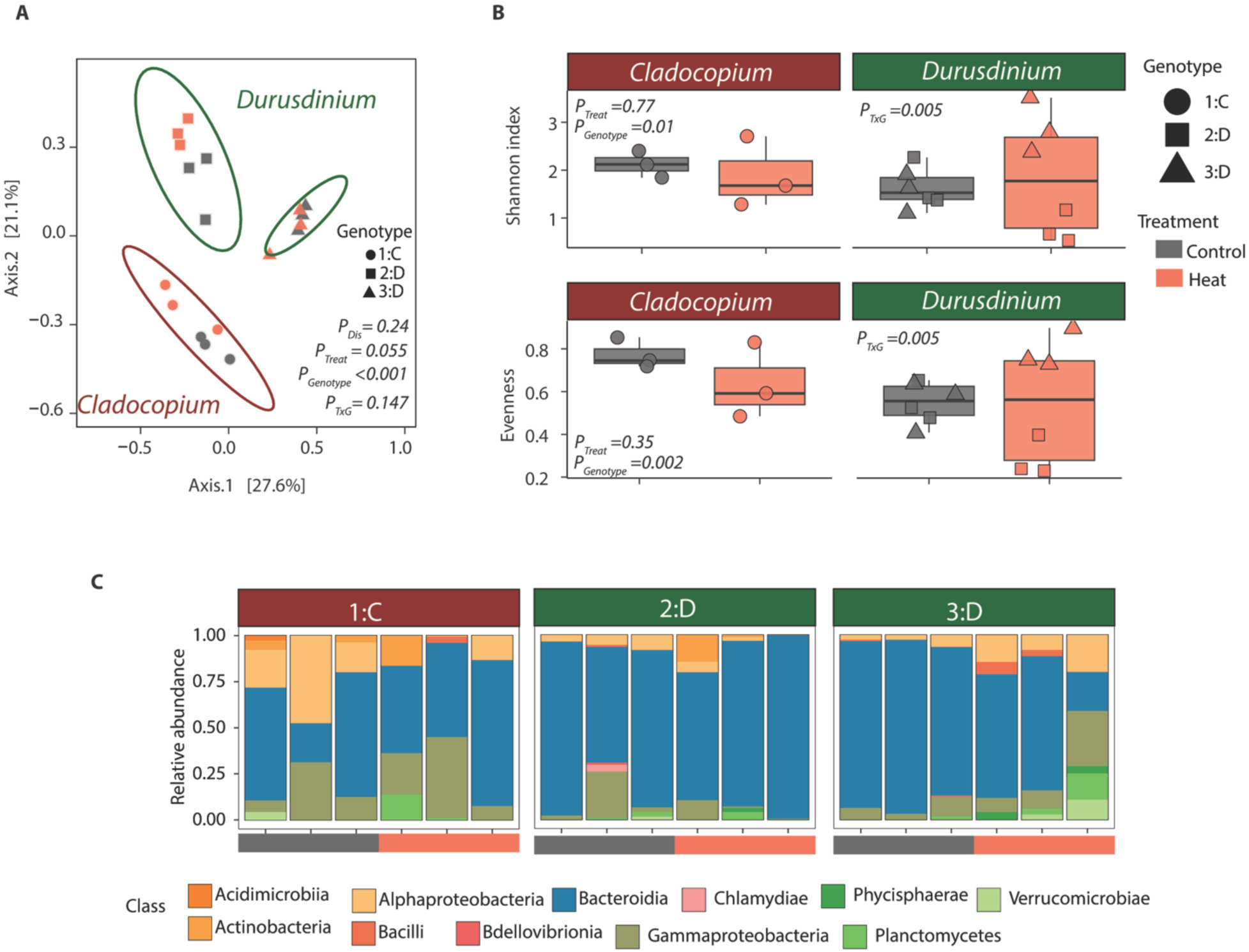
Impact of heat challenge on coral microbiomes. **(A**) Principal Coordinates Analysis (PCoA) of 16S rRNA gene sequencing data of microbiome communities based on Bray-Curtis dissimilarity, showing distinct clustering host genotype (P_Genotype_<0.001), but no significant clustering by heat treatment (P_Treat_=0.055, dispersion values across treatment or P_Dis_=0.24). (**B**) 16S Microbial alpha diversity metrics (Shannon index and evenness) between treatments and among host genotypes in *C-*hosting corals (left) and *D-*hosting corals (right). (**C**) Relative abundances of bacterial classes under heat challenge (orange) and control (grey) conditions separated by coral host-symbiont pairings.

Analysis of relative abundances showed that differences in microbiome communities were mainly driven by host:symbiont pairings at both the class (Fig. 6C) and family (fig. S7) levels. Genotype 1:C had a decreased abundance of Bacteroidia (mean decrease: 7.79 ± 1.82% SE, t(12) = 4.28, P = 0.0028, Fig. 6C, fig. S8A) and unchanged Gammaproteobacteria and Alphaproteobacteria under heat challenge (fig. S8, B and C). In contrast, *D*-hosting corals (specifically genotype 3:D), which showed the greatest tissue loss under heat treatment, experienced significant increases in relative abundance of Alphaproteobacteria (mean increase:13.35 ± 1.82% SE, t(12) = -7.338, p<0.0001, Fig. 6C, fig. S8C) and decreases of Bacteroidia (mean decrease: 35.52 ± 12.3% SE, t(12) = 2.89, P =0.03) under heat challenge (Fig. 6C; fig. S8A). These shifts were not observed in the other *D*-hosting genotype, 2:D. Our results suggest that although heat challenge induces shifts in microbiome communities in *C*- or *D*-hosting corals, host genotype largely determines microbiome restructuring and stability under thermal challenge in a laboratory setting.

### *D*-hosting corals experience greater tissue loss and lower survival than *C*-hosting corals under *Vibrio coralliilyticus* challenge

To determine whether *C-* or *D-*hosting *P. acuta* differed in their susceptibility to a pathogen, we exposed three host:symbiont pairings (1:C, 2:D, 3:D) to a known marine pathogen *Vibrio coralliilyticus,* and then compared tissue loss and survival outcomes. Tissue loss was strongly influenced by both pathogen treatment (ANOVA F=1754.86, df=1, p<0.001), and genotype (ANOVA F=443.03, df=2, p<0.001), and their interaction (ANOVA F=443.03, df=2, P<0.001, Fig. 7A). *D*-hosting corals experienced significantly greater tissue loss (75%) than their *C*-hosting counterparts (ANOVA p<0.001, Fig. 7A), with *C*-hosting corals not showing any evidence of tissue loss under pathogen treatment after seven days (Fig. 7A). Coral survival when infected with *V. coralliilyticus* was also significantly lower in both *D-*hosting genotypes relative to the *C*-hosting genotype (log-rank χ^2^=26.9, df=1, p<0.0001). Survival rates of *D-*hosting genotypes decreased at day 6 post infection, whereas no decrease in survival rate was observed in the *C*-hosting genotype (log-rank χ^2^=0, df=1 P=1). Together, these results suggest that algal type influences resistance to pathogen infection in the three host:symbiont pairings used in this study, at least in the case of *V. coralliilyticus*. Consistent with this pattern, reanalysis of lesion progression in corals exposed to Stony Coral Tissue Loss Disease (SCTLD)-infected *Diploria labyrinthiformis* revealed increased lesions in *D*-hosting corals compared with *C*-hosting corals (fig. S9, ANOVA P_Treatment_< 0.001), supporting a link between algal type and disease susceptibility in another coral species.

**Fig. 7.**
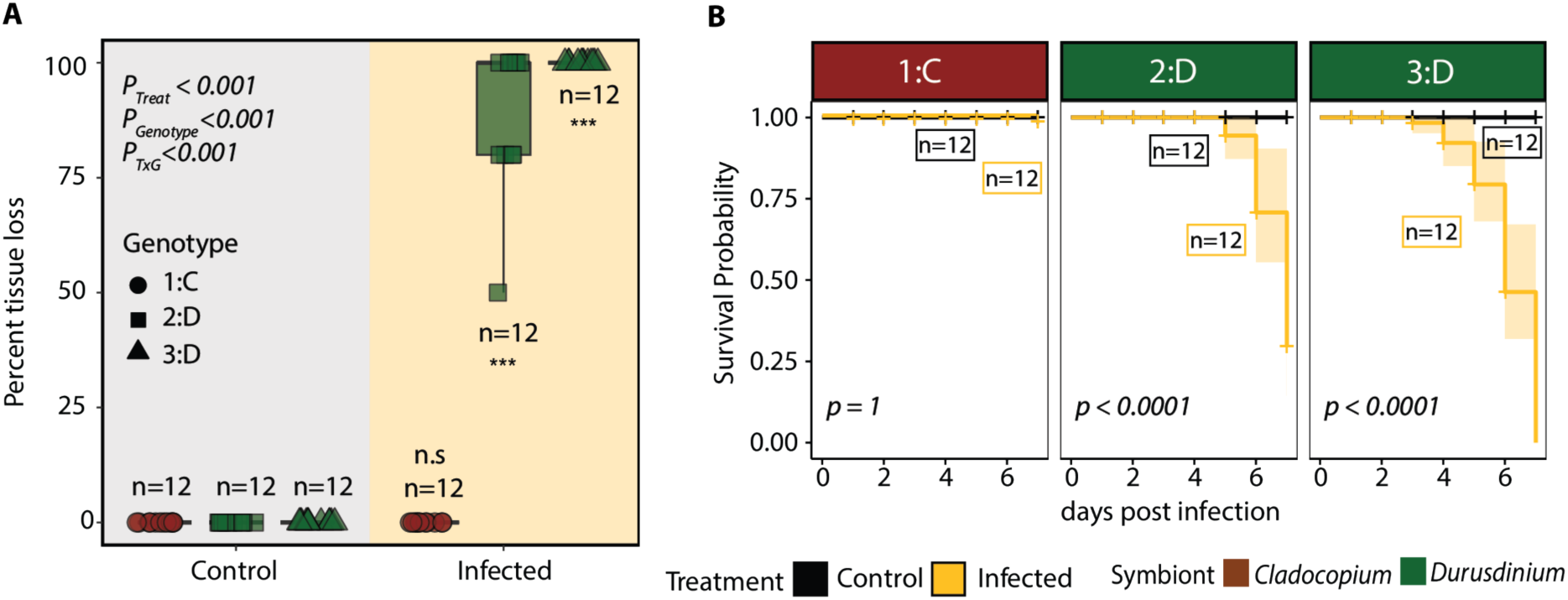
Algal type influences *P. acuta* coral tissue loss and survival following *Vibrio coralliilyticus* infection. (**A**) Percentage of tissue loss in *C*- (1:C) and *D-*hosting (2:D, 3:D) corals six days following *V. coralliilyticus* infection. *D-*hosting corals exhibited significantly greater tissue loss compared to the *C-*hosting coral (p<0.001). Symbols represent different genotypes, background color represents treatment (black = control with heat-killed *V. coralliilyticus*; yellow = infected with live *V. coralliilyticus*), box plot color represents algal symbiont type (brown = *C*-hosting, green = *D*-hosting). (**B**) Survival probability following six days of *V. coralliilyticus* infection. Both *D-*hosting genotypes infected with *V. coralliilyticus* had reduced survival (p<0.0001), while the *C-hosting* genotype exhibited no difference in survival between infected and control treatment. In both (**A**) and (**B**), n=12 individual replicate fragments per genotype per treatment. Error bars denote standard error.

### Enrichment of genes related to immunity and response to stress in other coral species when hosting *Durusdinium*

To establish whether the widespread enrichment of immune-related pathways in our *D*-hosting corals extends beyond our study system, we reanalyzed gene expression data from two independent studies that included *C-* and *D*-hosting corals (*32*, *33*). First, principal component analysis of adult *C*- and *D*-hosting *Pocillopora grandis* from (*32*) showed a clear separation of host gene expression profiles by algal symbiont type (Fig. 8A, P = 0.016). GO enrichment analysis of these data revealed an over-representation of *Biological Process* terms associated with immunity in *D*-hosting corals, which included immune system process [GO:0002376], DNA recombination [GO:0006310], and other functions such as ammonium transport [GO:0015696] (Fig. 8B). Next, we examined a dataset from (*33*) that experimentally infected *Acropora tenuis* recruits with *Cladocopium* or *Durusdinium*. Again, PCA indicated host gene expression separation by algal symbiont type (Fig. 8C, p<0.001). Notably, among enriched GO terms in *D*-hosting recruits were double-strand break repair (GO:0006302), cellular response to DNA damage (GO:0006974), and protein-containing complex organization (GO:0043933), suggesting heightened cellular surveillance and repair (Fig. 8D). Overall, these re-analyses confirm enrichment of immune and stress response as a conserved feature of *D*-hosting corals.

**Fig. 8.**
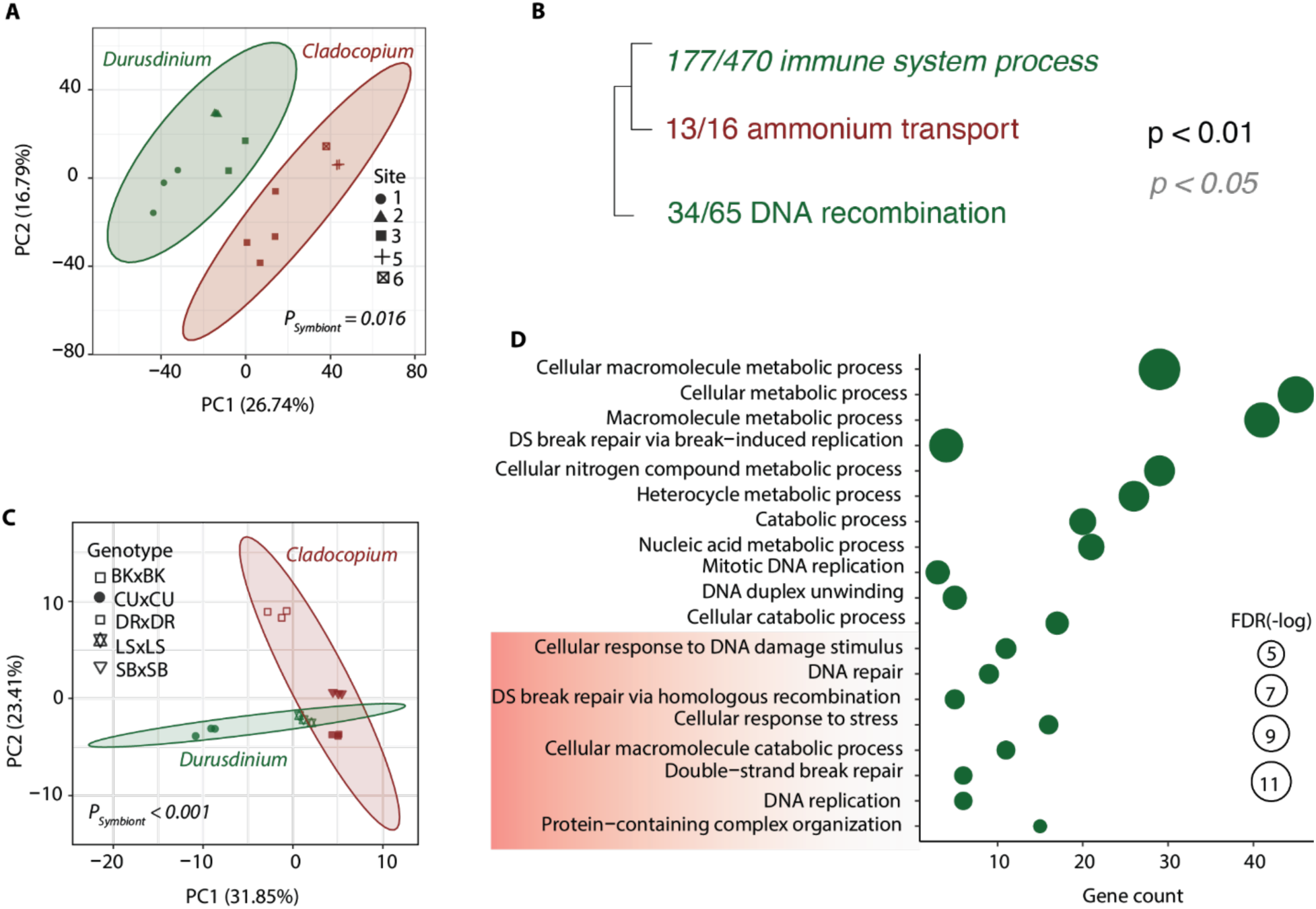
*D-*hosting corals consistently exhibit stronger immune and stress-related transcriptional responses relative to *C*-hosting corals across two previous studies. Data for (**A**) and (**B**) are from (*32*) and data for (**C**) and (**D**) are from (*33*). (**A**) Principal component analysis (PCA) of gene expression data from adult *Pocillopora grandis* hosting either *Cladocopium* (brown) or *Durusdinium* (dark green) collected from six reef sites (symbols). Axes are percent variation explained; *PERMANOVA* P_Symbiont_ = 0.016. (**B**) Gene Ontology enrichment of *C*- and *D*-hosting *P. grandis*, showing significant overrepresentation of immune system process genes in *D*-hosting corals, alongside other enriched categories (e.g., DNA recombination and ammonium transport). (**C**) PCA of gene expression of *Acropora tenuis* recruits infected with *Cladocopium* (brown) or *Durusdinium* (dark green) showing distinct expression profiles (*ADONIS* P_Symbiont_=.001). Individual host genotypes are shown by distinct symbols. Axes indicate percent variation explained. (**D**) TopGO enrichment showing upregulated genes in *D*-hosting recruits, highlighting the enrichment of cellular stress and DNA-damage response processes in *D*-hosting relative to *C-*hosting corals. Dot size denotes –log(FDR-corrected p-value).

## DISCUSSION

This study investigated how algal symbiont type affects baseline immunity, gene expression, and responses to multiple stressors in the tropical coral *P. acuta*. We demonstrate that algal type — *Cladocopium* vs. *Durusdinium* — alters *P. acuta*’s immunological status and its ability to respond to environmental and pathogenic stress by characterization of comparative physiology, host gene expression, and microbial communities. Moreover, hosting *Durusdinium* may induce a stressed state, as evidenced by increased expression of transcription factor NF-κB in *D*-hosting corals. These findings suggest that trade-offs exist when corals host different algal symbiont types and that the maintenance of symbiosis under environmental challenges is context-dependent.

### Higher baseline immune profiles in *D*-hosting than *C*-hosting corals

Gene expression profiling and Western blotting of NF-κB revealed distinct immunological profiles between *C*- and *D*-hosting corals at baseline. That is, *C*-hosting corals showed reduced expression of immune-related genes and NF-κB protein relative to *D*-hosting corals. Moreover, TNF receptor-associated factor (TRAFs), peroxidases, and members of the NF-κB pathway, all showed reduced expression in *C*-hosting corals, suggesting an internally entrenched immunological integration of the host when associated with *Cladocopium* algal symbionts (*13*, *34*). This association of immune suppression in the presence of *Cladocopium* in *P. acuta* could be an outcome of their coevolution (*25*), an association that may dampen host immune surveillance to ensure mutualism stability, which is a trait observed in other well-established and stable symbioses (e.g., legumes and squid) (*35*, *36*).

*Pocillopora acuta* hosting *Durusdinium* also had higher levels of NF-κB protein than *P. acuta* hosting *Cladocopium*. A similar pattern of NF-κB protein expression has been shown for the facultative anemone Aiptasia (*24*). These higher levels of NF-κB when hosting *Durusdinium*, perhaps as a stress-response factor, may act to frontload *P. acuta* and Aiptasia, and thus increase thermotolerance. NF-κB protein and mrRNA levels have also been shown to be upregulated by other stressors in Aiptasia (e.g., (*31*)). At first glance, this result could suggest a defensive or preemptive advantage against environmental stress; however, it is also possible that this heightened immune signature is due to the host identifying *Durusdinium* as a foreign symbiont (*30*). In general, literature spanning invertebrates and vertebrates suggests that mutualistic symbioses are typically dependent on the evasion or suppression of host immunity by the symbiont (*37*) or, conversely, an evolved symbiont tolerance by the host without induction of an inflammatory-type of response (*30*). In *P. acuta*, the failure of *Durusdinium* to fully evade or suppress host immune responses may reflect an incompatibility (*30*) or an association that is only recently tolerated (*38*).

In mammalian cells, the NF-κB p100 protein is in an inactive cytoplasmic state until it receives an upstream activating signal. NF-κB p100 is then activated by IKK-directed phosphorylation of three C-terminal serine residues, followed by proteasomal degradation of the C-terminal ANK repeat domain, which then allows for nuclear localization of the active, truncated NF-κB p52 (essentially RHD-only) protein (*39*). Our characterization of Pa-NF-κB showed that it has a linear structure—with an N-terminal RHD and C-terminal ANK repeats—that is similar to both mammalian NF-κB p100 and Aiptasia NF-κB. Nevertheless, our analysis revealed that there is a difference between *P. acuta* and Aiptasia NF-κB proteins. In *P. acuta*, the NF-κB protein is exclusively in its full-length form, whereas in Aiptasia, NF-κB is almost exclusively processed (*24*). Using human cell models, we have shown that the overexpressed Pa-NF-κB is largely in an inactive state, i.e., located in the cytoplasm in its full-length form, but enters the nucleus when the C-terminal ANK repeat domain is removed. Two pieces of evidence suggest that Pa-NF-κB ca be activated by C-terminal processing: 1) there is some constitutively processed (∼50 kDa) Pa-NF-κB in transfected HEK 293T cells (Fig. 2C and D), and 2) expression of a constitutively active form of human IKKβ can result in increased processing of Pa-NF-κB (fig. S3C), presumably by phosphorylating C-terminal serine residues. Essentially identical findings have been made with Aiptasia and *Orbicella faveolata* NF-κB proteins in human cells (*13*, *29*). Why there is a difference between the Aiptasia and *P. acuta* NF-κB proteins in their native organisms—Ap-NF-κB processed vs Pa-NF-κB non-processed—is not known. Still, it is intriguing that the NF-κB protein is expressed at higher levels in both Aiptasia and *P. acuta* when they are hosting the thermotolerance-inducing algal symbiont *Durusdinium* versus *Cladocopium* (*24*).

Maintaining constitutive higher immunity when hosting *Durusdinium* algae may also have detrimental “trade-off” consequences (*40*). For example, continued activation of gene pathways such as NF-κB, TNF, and MAPK in *D*-hosting cnidarians may drive inflammatory injury (*41*), induce apoptosis (*42*), and alter tissue homeostasis (*43*), any one of which can become deleterious under heat or pathogen challenge. Importantly, the elevated expression of immune-related genes and higher NF-κB levels in *D*-hosting corals mirrors phenomena experienced during dysbiosis (*44*) and symbiont shuffling (*38*), because incompatibility between hosts and symbionts leads to elevated immune response and disruption of homeostasis (*30*). Supporting this interpretation, reanalysis of *P. grandis* transcriptomes revealed enrichment of immune activation gene sets in *D*-hosting relative to *C*-hosting corals. Likewise, coral recruits experimentally infected with *Durusdinium* showed gene expression patterns consistent with cellular stress and immune activation, suggesting that *Durusdinium* association is more broadly perceived by the host as foreign or stressful across species and life stages. These observations align with our hypothesis that heightened immunity in *D*-hosting corals is a less adaptive association, rather than an effective mutualistic integration. Such immune activation is often energetically costly for the host (*45*), suggesting that the heightened immune and/or stressed state in *D*-hosting corals likely comes at a cost. For example, acute infection with an opportunistic pathogen typically causes inflammation, with cascading effects on metabolism (e.g., hypermetabolism to support immune activation) and nutrition (*45*). Together, these findings suggest that immune suppression is a feature of symbiotic integration with *C*-hosting corals being an optimized low-cost association, in contrast to the energetically expensive partnership in *D*-hosting corals. While we were not able to fully disentangle host genotype and algae type in this study, future work on reef-building coral species where algal manipulation has been achieved (*Galaxea fascicularis* (*46*); *Diploastrea heliopora, Dipsastraea pallida, Echinopora lamellosa, Platygyra daedalea, Porites lobata* and *Stylophora pistillata* (*47*)) represent a promising area of research.

### *C*- and *D*-hosting corals exhibit different tissue responses to thermal stress

In their hosts, *Durusdinium* has been widely considered to be an inducer of tolerance to stressors such as heat (*48*). However, our results indicate that the short-term benefit for thermotolerance of hosting *Durusdinium* comes at a cost. Here, the *C*-hosting *P. acuta* genotype experienced greater reductions in algal density and photochemical efficiency under heat challenge than the two *D*-hosting genotypes, consistent with the known sensitivity of *C*-hosting corals to thermal stress (*49*). However, the bleached fragments from *C*-hosting *P. acuta* were able to maintain tissue integrity. In contrast, despite retaining relatively efficient photophysiology, *D*-hosting genotypes experienced greater tissue loss than the *C*-hosting genotype. Similar patterns have been documented in *M. capitata* and *Galaxea fascicularis*, where *D*-hosting corals experienced severe tissue loss and mortality while *C*-hosting corals bleached under prolonged stress exposure (*19*, *49*), suggesting that this trade-off extends beyond *P. acuta*. These findings initially seem contradictory given the enhanced survival of *D*-hosting corals during *in situ* thermal disturbances (*50*, *51*). Yet this photophysiological resistance likely confers their field survival advantage. Natural heat wave events may not have reached the duration or magnitude necessary to induce tissue loss (here, 32°C for six days), though intensifying marine heat waves may increasingly exceed this buffering capacity (*52*). In addition, the shared transcriptomic signatures observed here—elevated baseline immunity, chronic inflammatory signaling under heat stress, and metabolic dysregulation in *D-*hosting genotypes of *P. acuta*—also was observed in our reanalysis of *D*-hosting *P. grandis* and *A. tenuis* recruits experimentally infected with *Durusdinium*. These signatures suggest shared physiological costs that can manifest as tissue loss under extreme or prolonged thermal stress across diverse coral-*Durusdinium* associations.

Persistent induction of the TNF pathway in *D*-hosting corals under heat challenge potentially links chronic inflammatory signaling to tissue loss. *D*-hosting corals showed elevated expression of multiple TRAF homologs (*TRAF2*, *TRAF3.1*, *TRAF3.2*, *TRAF3.4*; TNF signaling adaptors), caspase domain (apoptotic effectors), *BCL3* (NF-κB-associated protein), *JUN* (stress-response transcription factor), and *IFIH1* (viral RNA sensor triggering interferon responses), under thermal challenge. This coordinated upregulation of TNF pathway components and inflammatory mediators indicates activation of a pro-inflammatory transcriptional program characteristic of immune activation and tissue damage responses. Among the *P. acuta TRAF3* homologs, three were significantly upregulated in *D*-hosting corals under heat challenge, but it is worth noting that other *TRAF3* homologs showed variable expression patterns, reflecting the complexity of having multiple copies of gene families in corals (*53*). Persistent activation of TNF pathway signaling drives inflammatory cascades that promote apoptosis (*42*), tissue injury (*41*), and homeostatic disruption (*43*) which becomes detrimental under prolonged stress. This inflammatory signature is consistent with patterns observed in other heat-stressed corals, where upregulation of TNF signaling and apoptotic pathways has been linked to tissue damage and symbiosis breakdown (*42*). Together with downregulation of the tissue repair factor FGF18 (fibroblast growth factor for tissue repair), these expression patterns support a link between inflammatory activation and tissue loss in *D*-hosting corals. These patterns reflect a trade-off, where hosting *Durusdinium* ensures photophysiology resistance, but at the expense of host structural integrity (*17*, *54*), possibly due to higher constitutive host immunity derived from host-symbiont incompatibility.

Under elevated temperatures, *D*-hosting corals also exhibited higher expression of genes associated with the MAPK pathway relative to *C*-hosting corals, and this pathway is essential in stimulated cellular processes (*55*) and symbiosis (*30*). While corals hosting either symbiont type downregulated expression of some genes encoding key signaling components–-such as *Myd88* (TLR signal adaptor), *FGF18* (fibroblast growth factor for tissue repair), and *FGFR4* (FGF receptor tyrosine kinase)–under heat challenge, only *D*-hosting corals upregulated certain effector genes in the MAPK pathway. These MAPK pathway genes included *PLA2G4A* (phospholipase producing inflammatory lipid mediators), *MAP3K20* (stress-activated MAP kinase kinase kinase essential for algal symbiont take up), and *MKNK1* (MAPK-regulated translation modulator), which reflects a bias towards a pro-inflammatory and stress-amplifying response. This selective activation was also accompanied by downregulation of *gal_lectin* (symbiont recognition lectin, (*56*)) in *D*-hosting corals, which suggests the emergence of symbiotic incompatibility (*57*).

Elevated NF-κB signaling in *D*-hosting corals under heat challenge suggested chronic immune activation as a potential driver of tissue deterioration. Coral genotypes hosting either symbiont type showed increased expression of transcripts for *NFKB1* (transcriptional regulator of inflammation), NF-κB-associated protein *BCL3*, and *TRAF3* under heat challenge; however, the magnitude of upregulation was significantly greater in *D*-hosting than *C*-hosting corals. Additionally, there was an upregulation of *CSNK1G*2 (casein kinase that modulates NF-κB activity). Overall, these results suggest sustained NF-κB upregulation in *D*-hosting corals. This pattern of gene expression is consistent with prolonged expression of inflammatory mediators in *D-*hosting corals, which has been observed in vertebrate models wherein chronic NF-κB activation predisposes animals to immunopathology (*58*). The concurrent induction of apoptotic markers such as apoptosis effectors (*CASP7*), viral response genes (*IFIH1* a viral RNA sensor triggering interferon responses), and mitochondrial dysfunction markers (*DNM1L*: dynamin-related protein mediating mitochondrial fission) also resembles mammalian sterile inflammation, in which endogenous stimuli (in this case, symbiont stress signals) provoke excessive immune activation (*59*), leading to tissue loss (*60*). While acute stimulation of these pathways may be beneficial (*61*), elevated or dysregulated immune stimulation could result in cellular damage and compromised tissue integrity if experienced over extended time periods (*62*).

Metabolic reprogramming also signals host-algal partnership destabilization (*63*). Symbiosis-related genes such as *amt-1* (ammonium transporter) and *NPC2* (cholesterol transporter) remained unchanged in *C*-hosting corals under heat challenge, but were downregulated in *D*-hosting corals, implying disrupted nutrient transfer (*31*). Reduced assimilation of ammonium in the host cell supports greater nitrogen transfer to the symbiont, potentially disrupting symbiont nitrogen-limitation, a defining feature of stable coral-symbiont partnerships (*64*). However, only *D*-hosting corals induced aldehyde dehydrogenases (*ALDH1L2*, *ALDH3B1*) and *ALOX5* (arachidonate lipoxygenase), which are genes associated with lipid peroxidation and aldehyde dehydrogenase involved in lipid-based carbon acquisition (*65*). These patterns suggest that there is greater metabolic disruption in *D*-hosting corals under heat challenge, which then contributes to symbiosis instability.

We also investigated whether different stress responses were associated with differences in microbiome compositions between *C*- and *D*-hosting corals. Overall bacterial communities were shaped by host-symbiont pairings, with limited shifts in response to elevated temperature in both *C*-hosting and *D*-hosting corals. In *C*-hosting corals, bacterial taxa such as Gammaproteobacteria and Bacteroidia remained stable under heat stress, whereas nitrogen-fixing and antioxidant producing Alphaproteobacteria decreased– a pattern also observed during polyp bail-out in stressed corals (*66*). Loss of nitrogen-fixing and antioxidant-producing taxa has been associated with reduced nitrogen availability and weakened oxidative-stress responses in bleaching corals (*67*), though whether these microbial shifts drive or simply accompany physiological decline remains unresolved. In contrast, one *D*-hosting coral (3:D) exhibited a decrease in Bacteroidia and an increase of opportunistic Alphaproteobacteria, which could be attributed to epithelial damage and immunological hyperactivation. Other genotypes maintained stable communities, with genotype-specific susceptibility to microbiome destabilization and impaired nutrient exchange. Together, our findings reveal a holobiont-level trade-off in coral thermal tolerance, where *D*-hosting corals maintain photosynthetic function but suffer tissue loss coupled with immune, metabolic, and microbial dysregulation. In contrast, *C*-hosting corals bleached following a decline in nitrogen-fixing and antioxidant-producing bacteria yet retained structural tissue integrity–demonstrating that symbiont stress tolerance does not guarantee holobiont survival. While these findings reveal distinct thermal stress strategies, they are based on one *C*-hosting versus two *D*-hosting corals, and algal symbionts were only characterized at the genus level. Given that ITS2 sequencing can fail to capture finer-scale genetic diversity within *Cladocopium* and *Durusdinium* (*68*), future studies with additional host genotypes and algal strains coupled with functional microbial analyses (e.g., metatranscriptomics, metabolomics) may be required to establish causal mechanisms linking symbiotic partnerships to holobiont performance.

### Association with *Durusdinium* algal symbionts impairs pathogen responses

To determine whether the higher baseline immune status of *D*-hosting corals would confer pathogen tolerance, we conducted a pathogen challenge experiment using the natural bacterial pathogen *V. coralliilyticus*. Our results showed that despite higher baseline immune gene expression and NF-κB protein levels relative to the *C-*hosting coral, both *D*-hosting corals experienced extensive tissue loss and lower survival following infection with *V. coralliilyticus*. In contrast, the *C*-hosting coral, which exhibited a comparatively muted immune expression profile, exhibited no tissue loss and high survival probability. This outcome contradicts the assumption that high baseline immunity confers protection against pathogens (*69*). Instead, we show that immune stimulation in *D*-hosting corals reflects a chronic, dysregulated inflammatory state rather than an adaptive, protective immune readiness. Such persistent immune stimulation may arise as a physiological cost of maintaining symbiosis with a less compatible algal partner (*30*, *38*), ultimately impairing the host’s ability to mount an effective response when challenged by pathogens.

Immune exhaustion, which is a condition in which constitutive or chronic stimulation of immune mechanisms is followed by diminishing responsiveness upon new infection (*70*), may explain the patterns observed in *D*-hosting corals. In this case, the host is already initiating an immune response as it goes on to monitor and manage less compatible algal symbiont relationships, thus limiting its capacity to resist pathogenic infection. Immune misdirection (*71*) may also explain these patterns, where immune stimulation in *D*-hosting corals becomes misdirected or biased against foreign symbiont control at the cost of pathogen defense, leading to ineffective protection. Whether through exhaustion or misdirection, these immune dysregulation mechanisms link constitutive immune activation in *D*-hosting corals to the impaired pathogen resistance and tissue loss that we observed under thermal challenge. Together, these findings suggest that algal symbioses influence coral host thermotolerance, with corals associating with *Cladocopium* favoring long-term host survival and corals hosting *Durusdinium* prioritizing short-term stress management at the cost of tissue integrity and immune function.

By integrating data across scales, this study provides a mechanistic framework for understanding the trade-offs of hosting *Cladocopium* versus *Durusdinium* under thermal challenge. The conservation of gene expression signatures across multiple coral species hosting *Durusdinium* (*P. acuta*, *P. grandis*, *A. tenuis*) suggests that these immune costs represent a fundamental constraint of these host-symbiont pairings. While shuffling to thermotolerant symbionts may offer short-term benefits under acute thermal stress, the long-term consequences of chronic immune activation, pathogen susceptibility, and tissue degradation could undermine coral survival in multi-stressor or extreme heat environments. This hypothesis is supported by recent observations of severe tissue loss and mortality in *D*-hosting corals under prolonged heat (*19*, *49*) and by *D*-hosting corals showing elevated lesion progression rates during disease exposure (*54*). It is important to note that this study only used one *C*-hosting and two *D*-hosting corals for all of the challenge assays, and host genotype has been shown to play a significant role in coral stress responses (*9*).

## MATERIALS AND METHODS

### Identification of host and algal type pairs

Multiple colonies representing five distinct *P. acuta* genotypes acquired from the aquarium trade have been independently maintained at Boston University for more than nine years under common garden conditions (temperature: 26 ± 1 °C, salinity: 34-35 ppt, minimum photosynthetic photon flux density:120 µmol m⁻² s⁻¹ on an 8:16 hour Light:Dark cycle). Corals were fed three times a week with newly hatched brine shrimp (*Artemia salina*). These coral genotypes (fig S10A) naturally predominantly associate with one of two Symbiodiniaceae genera: *Cladocopium* and *Durusdinium* (formerly Clade C and D, respectively) and are referred to as *C*- and *D*-hosting corals throughout the manuscript (fig. S10B and S11). All five host-symbiont pairings, denoted using the format *Genotype:Algal Symbiont Type* (1:C, 2:D, 3:D, 4:C, 5:D), were included in baseline gene expression comparisons. Due to coral husbandry challenges, all follow-up experiments and measurements were conducted in genotypes 1:C, 2:D and 3:D. Details on specific genotype inclusion and replication are explained in each relevant methods subsection.

### Coral host genotype and dominant algal symbiont assessments

Total RNA was isolated from homogenized tissue of three independent fragments per colony for all five host-symbiont pairings (1:C, 2:D, 3:D, 4:C, 5:D; N = 15 total samples) using the RNAqueous Total RNA Isolation Kit (Invitrogen) following the manufacturer’s protocol. An additional DNA removal step was performed using DNA-free™ DNA Removal Kit (Invitrogen). TagSeq library preparation and sequencing were performed in two batches. The first batch (genotypes 1:C, 2:D, and 3:D; N = 9 samples) was processed at the University of Texas at Austin Genome Sequencing and Analysis Facility (GSAF) on the NovaSeq6000 platform. The second batch (genotypes 4:C and 5:D; N = 6 samples) was processed at Tufts University Core Facility (TUCF) Genomics on the Illumina HiSeq 2500 platform. Identification of coral host clones was conducted as described by Rivera et al. (*21*). Briefly, cleaned reads were mapped to the concatenated genome of *P. a acuta* (*72*), and an algal symbiont database (transcriptomes of *Symbiodinium spp*, *Breviolum spp*, *Cladocopium spp*, *Durusdinium spp*) following (*73*) using Bowtie2 v2.5.1 (*74*). Reads mapping to *P. a acuta* were then used to detect single nucleotide polymorphisms (SNPs) using ANGSD v0.935 (*75*). A hierarchical clustering tree (hclust) of samples based on pairwise identity by state distances calculated in ANGSD determined genetic similarity, which facilitated the identification of host clones (fig. S10A). To characterize the relative proportions of reads belonging to each algal symbiont genera in each library, reads mapping to the algal symbiont database were used to calculate the relative abundances of transcripts mapping to each algal genus. All samples had 98.1 ± 0.017 % (Mean algal transcript fraction ± SD) of reads mapping to a single genus, which allowed for the identification of *Cladocopium*- and *Durusdinium*-dominated individuals (fig. S10B).

Genomic DNA (gDNA) was retrieved from RNA isolations prior to the DNA removal step for one representative colony per genotype across all five host-symbiont pairs (1:C, 2:D, 3:D, 4:C, 5:D; N = 5 total samples). The ITS2 region of Symbiodiniaceae ribosomal DNA (*76*) was amplified following standard protocols (see Supplemental Methods) and sequenced (250 bp paired-end reads) at the TUCF on an Illumina MiSeq 2500. ITS2 data were submitted to SymPortal (*77*) to characterize ITS2 type profiles. Relative abundances of majority ITS2 sequences were visualized using phyloseq (*78*).

### Baseline gene expression profiling of *C*- and *D*-hosting corals

Raw reads were processed with the TagSeq pipeline (https://github.com/z0on/tag-based_RNAseq). Briefly, Fastx_toolkit trimmed adapters and poly(A)+ tails, and sequences shorter than 20 base pairs were removed. Reads that passed quality filtering were then mapped to concatenated genomes of *P. acuta* (host genome, (*72*)), *D. trenchii* ((*53*)), and *C. goreaui* ((*79*)) with Bowtie2 v2.5.1 (*74*). Reads mapping to *P. acuta* were used in downstream analyses. Baseline gene expression profiling included three replicates of all five genotypes: 1:C, 4:C, 2:D, 3:D, and 5:D. To account for sequencing run batch effects, ComBat-seq was used to model out the effects of sequencing batch prior to differential expression analysis (*80*). Differential gene expression analysis of *C*- vs *D*-hosting corals was performed using DESeq2 v1.44.0 (*81*) with a design formula of ∼ Genotype (FDR <0.05). Principal Component Analysis (PCA) was conducted to characterize overall variation in gene expression among *C*- and *D*-hosting corals. A PERMANOVA tested for significant differences in clustering using the adonis2 function in vegan (*82*). All tracking statistics of reads are presented in table S14. Raw sequence reads can be found on the NCBI short read archive (SRA) database under BioProject accession number PRJNA1314775. All data and code can be found at https://github.com/jpdaanoy2024/Proj_Pacuta_Immuno.

Gene Ontology (GO) enrichment analysis was conducted on ranked p-values from *DESeq2* testing for differences between *C*- and *D*-hosting corals using Mann-Whitney U tests (*83*). *Biological Process* categories of GO terms were shown as clustered dendrograms according to shared genes among GO terms. A broad enrichment of immune process was identified. The list of immune-related genes from (*30*) were then manually retrieved from the significant GO terms and a heatmap was created using the pheatmap package (*84*).

### Western blotting of NF-κB proteins in *C*- vs *D*-hosting corals

Protein isolation from *P. acuta* tissue and Western blotting were performed as described previously (*13*). Coral tissue from four fragments from three host-symbiont pairings (1:C, 2:D, 3:D; N=12 total samples) were homogenized in 100 µl of 2X SDS sample buffer (0.125 M Tris-HCl, pH 6.8, 4.6% w/v SDS, 20% w/v glycerol, 10% v/v β-mercaptoethanol, 0.2% w/v bromophenol blue) with a mortar and pestle, and samples were then incubated at 95°C for 10 min. Homogenates were centrifuged at 13,000 rpm for 10 min, and the supernatants were collected. For electrophoresis, equal amounts of total protein from each sample were separated on a 7.5% SDS-polyacrylamide gel and transferred to a nitrocellulose membrane. The membrane was blocked for 1 h in TBST (10 mM Tris-HCl [pH 7.4], 150 mM NaCl, 0.05% v/v Tween 20) with 5% non-fat powdered milk, then incubated overnight at 4°C with primary rabbit anti-Ap-NF-κB antiserum (1:5000) (*13*). After five TBST washes, the membrane was incubated for 60 min in HRP-conjugated goat anti-rabbit secondary antibody (Cell Signaling Technology, Cat. # 7074V) at a 1:4000 dilution. After five TBST and two TBS washes, immunoreactive proteins were detected using the SuperSignal West Dura Extended Duration Substrate (Fisher Scientific) and were imaged using a Sapphire Biomolecular Imager. Pa-NF-κB band intensities were quantified in each lane using ImageJ, with Ponceau S staining serving as a total protein loading control. Relative Pa-NF-κB levels were calculated as the ratio of Pa-NF-κB protein to Ponceau S staining for each sample, which were themnormalized to the average of the extracts from *C*-hosting tissue (set to 1.0). Differences in Pa-NF-κB levels across host-symbiont pairings were tested using an ANOVA.

### Characterization of the *P. acuta* transcription factor NF-κB

The *P. acuta* NF-κB protein was characterized using methods that we have previously used for other basal NF-κB proteins (*24*, *29*). We searched the annotation list associated with the *P. acuta* genome for genes annotated as NF-κB, and identified a single NF-κB protein coding sequence. To analyze functional domains of the Pa-NF-κB protein we subcloned human codon-optimized cDNA sequences (GenScript; fig. S2B) into expression vectors for FLAG-tagged full-length Pa-NF-κB (aa 2-919), an RHD-GRR truncated protein (aa 2-461), and a MYC-tagged C-terminal ANK repeat-containing Pa-Cterm protein (aa 461-919) (Fig. 2B). Expression of these proteins was analyzed by transfection of 293T cells, followed two days later by anti-FLAG or anti-MYC Western blotting of cell extracts (*85*). As a negative control, we analyzed extracts from cells transfected with the empty vector, and as a positive control, we expressed a FLAG-tagged NF-κB protein from the sea anemone *Nematostella vectensis*, which is a naturally truncated RHD-GRR protein that we have characterized previously (*27*). To assess the DNA-binding activity of these proteins, we used extracts from the transfected 293T cells in an ELISA-based κB site-binding assay that we recently developed (*28*).

For analysis of subcellular localization of the Pa-NF-κB proteins, we transfected chicken DF-1 fibroblasts were transfected with our epitope-tagged expression vectors Two days later, cells were fixed in 100% methanol for 10 min at -20°C, then subjected to indirect immunofluorescence using anti-FLAG or anti-MYC primary antiserum as described previously (*85*). Stained cell samples were excited at 480 nm (FLAG) or 540 nm (MYC) and emission was detected at 525 nm (FLAG) or 600 nm (MYC) under 40X magnification on an Olympus BH-2 binocular microscope. To determine whether the Pa-RHD and Pa-Cterm repeat sequences can directly interact, we co-transfected 293T cells in a 100-mm tissue culture plate using polyethylenimine with 5 μg of the FLAG-tagged Pa-RHD plasmid or the empty vector and with 5 μg of the MYC-tagged Pa-Cterm plasmid or the empty vector (see fig. S3B), as described previously(*28*). Two days later, cells were lysed and we performed an anti-FLAG bead immunoprecipitation, followed by Western blotting (*85*).

For analysis of in vivo processing of Pa-NF-κB, we co-transfected 293T cells with equal amounts of plasmids for FLAG-Pa-NF-κB and a constitutively active HA-tagged form of human IKKβ (IKKβ-SS/EE) or the empty vector. Two days later, cell extracts were analyzed by Western blotting (*13*) with either anti-HA or anti-FLAG antiserum, and the amount of processed (∼50 kDa) Pa-NF-κB was compared to the amount of processed plus full-length Pa-NF-κB (fig. S3C). Finally, we used PyMOL (*86*) to compare the AlphaFold3-predicted structure of the Pa-RHD dimer on a consensus double-stranded NF-κB site (5’GGGAATTCCC3’) to the structures of mouse NF-κB p50 and Nv-NF-κB on the same site, as we have described previously (*28*) (see fig. S12 for the RHD sequences used for modeling).

### Thermal challenge mesocosm experiment

Thermal challenge mesocosm experiments and subsequent physiological and molecular measurements were conducted on three *P. acuta* host-symbiont pairings (1:C, 2:D and 3:D) in 40 L tanks supplied with filtered seawater. Six independent fragments from three host-algal symbiont pairs (1:C, 2:D, 3:D; N=18 total fragments) were randomly assigned to one of three experimental tanks set to 27 °C (control) or one of three heat challenge tanks (temperature increased 1°C day−1 from 27°C to 32°C, then maintained at 32°C for six days). Three replicate tanks were used for each temperature treatment, and temperatures were maintained by Aqualogic Digital Temperature Controllers connected with aquarium heaters for precise control of water temperatures. Each tank contained one fragment from each of the three host-symbiont pairings. Fragments were rotated throughout the experiment to avoid potential differences in light exposure and water flow within the tanks. Temperature and salinity were monitored three times daily using a YSI pro30 multiprobe. All setups were illuminated under low photosynthetic photon flux density (∼80 µmol m−2 s−1) on a 7.5:16.5 h L:D cycle. All samples were flash-frozen in liquid nitrogen on the final experimental day and stored at -80 °C.

To assess symbiont performance, algal symbiont photochemical efficiency of Photosystem II (Fv/Fm) was measured daily throughout the experiment using a Walz Junior-PAM™. To estimate changes in red channel intensity, a proxy for bleaching (*87*), photos of all fragments were taken on Day 0 and Day 12 on an iPhone 13. Photos were white-balanced in Photoshop v25.1 and uploaded to MATLAB for color intensity analysis following Winters *et al.*(*87*). Ten random points on each coral image were selected, and red channel intensities were quantified and averaged for each fragment. To account for baseline differences in red channel intensity among individual fragments, we calculated the relative change in red channel intensity as (Day 12 - Day 0) / Day 0. To estimate algal density from coral fragments at the end of the experiment, coral tissue was removed from skeletons via airbrushing using an airgun with sterile saltwater. The symbiont pellet was resuspended in 1 ml of sterile seawater. Symbiont densities (cells/cm^2^) were quantified for three replicate 10 μl tissue slurry aliquots per fragment using a hemocytometer under a light microscope. Counts were averaged and normalized to live coral tissue surface area. To quantify total symbiont chlorophyll (*a* + *c*; μg/cm²), symbiont tissue extracts were centrifuged, and resulting pellets were left overnight at 4°C in 1 ml of 90% acetone in the dark. Sample absorbances at 630 and 663 nm were measured for three 200 μl aliquots of each sample on a Synergy H1™ Microplate Reader spectrophotometer (see Supplemental Methods).

Tissue surface area calculations were conducted by 3D scanning coral skeletal fragments using an EIN-SCAN SE with the HDR and texture scanning options enabled. While total host protein (mg/cm²) was determined using a Bradford assay with Bovine Serum Albumin (BSA) standards (see Supplemental Methods).

Daily tissue loss was evaluated through visual inspection by the same researcher to minimize inter-observer variability. Percentage tissue loss was based on tissue loss relative to overall live tissue prior to heat treatment. Tissue loss was defined as the emergence of dead or necrotizing tissue or exposed skeleton with sloughing tissue. Daily photographs were not conducted to reduce handling disturbance during experimental treatments.

Physiological trait data (F_v_/F_m_, algal density, red channel intensity, total chlorophyll, total protein, tissue loss) from the final experimental time point were analyzed using linear mixed-effects models with the structure: Dependent variable ∼ Treatment * Genotype + (1|tank), where Treatment represents control vs. heat challenge, Genotype represents the unique host-algal genotype pairings (1:C, 2:D, 3:D), and tank accounts for variation among experimental tanks within a treatment. Analyses were performed on raw values except for algal density, which was normalized relative to control means (by genotype) due to baseline differences between *C*-hosting and *D*-hosting corals. For traits where interaction of temperature treatment and genotype showed a significant (p < 0.05) effect, Tukey’s HSD *post hoc* tests independently compared treatment responses for each genotype. For time-series data (F_v_/F_m_ , tissue loss), we used linear mixed-effects models with fixed effects of temperature treatment, genotype, and day, and a random effect of tank. Tukey’s HSD *post hoc* analyses were conducted when main effects of treatment and genotype were detected. Data were normalized to genotype-specific control means to visualize treatment effects. Additional details on statistical results for all physiological trait data, including model structures, sample sizes, test statistics, degrees of freedom, and p-values, are reported in Supplementary Information. All statistical analyses and data visualizations presented here were performed using R v4.4.0 (*88*).

### Gene expression profiling of *C*-and *D*-hosting corals under heat stress

Total RNA of corals exposed to heat treatment was extracted and prepared for sequencing as described above with baseline and heat treatment libraries of 1:C, 2:D and 3:D being sequenced at the same time. Three replicates from each of the three host-symbiont pairings (1:C, 2:D and 3:D) in heat treatment tanks (∼600 ng per sample, total of 9 samples) were sent to University of Texas at Austin GSAF for TagSeq library preparation and sequencing (single-end, 100 bp) on a NovaSeq6000 platform. Quality trimming, host count generation, and outlier detection were performed following methods outlined above and all analyses described here were performed on baseline and heat-treated samples together (N=18). No outliers were detected. To account for variation attributable to host genotype (*9*), which is confounded with algal symbiont type in our experimental design, we applied ComBat-seq (*80*), as the covariate and temperature as the variable of interest (Fig. S13A–C). This correction removed host genotype-associated variation while preserving temperature-associated biological variation among corals hosting different algal types. Trimmed, batch-corrected data were then included in a DESeq2 (*81*) model, which accounted for the main effect of temperature treatment. DEGs for Heat vs Control were determined using an FDR-adjusted p-value of <0.05. Expression data were *vst*-normalized and a Principal Component Analysis (PCA) was conducted to characterize variation in gene expression between treatments. To confirm that this normalization approach did not influence our conclusions, we performed parallel differential expression analyses on non-batch-corrected data. These included: (a) a DESeq2 model accounting for unique genotype pairings (∼ *Genotype + Treatment + Genotype:Treatment*) to capture holobiont-specific responses to heat treatment, followed by PCA to assess overall expression patterns between *C*-hosting and *D*-hosting genotypes (fig. S14); and (b) separate analyses within each algal symbiont type to assess treatment effects (*C*-hosting model: ∼Treatment; *D*-hosting model: ∼Treatment + Genotype). All analyses yielded consistent results (fig. S15A–B, S16A–B). All tracking statistics of reads are presented in table S14. Sequence reads can be found in the NCBI short read archive (SRA) database under BioProject accession number PRJNA1314775.

Weighted Gene Correlation Network Analysis (WGCNA; (*89*)) was performed on all genes whose basemean values were >3. Data were *vst*-normalized using *DESeq2* (*81*). Outlier samples were checked within the WGCNA package, and no outliers were detected. The “pickSoftThreshold” function explored soft thresholds from 1 to 30 and a value of 18 was chosen, which corresponded to a signed R^2^ >0.90. Signed connectivity among genes was determined, and eigengene expression of these modules was correlated to temperature treatments and physiological traits (minModuleSize=95, MEDissThres= 0.25). GO enrichment analysis of each module that was highly correlated to heated treatments was performed as described above with the modification of using Fisher’s exact tests (presence/absence in a module) instead of continuous ranked p-values. We then manually searched modules associated with heat treatments for genes annotated as Tumor Necrosis Factor (TNF), MAPK, and NF-κB (*30*), and symbiosis/stress genes (*31*) in the *P. acuta* genome, which were visualized in the pheatmap package in R.

### Microbiome profiling of *C-* and *D*-hosting corals under thermal challenge

Genomic DNA was co-isolated from RNA isolations by taking an aliquot from the extraction prior to DNA removal, following protocols validated in previous coral microbiome studies (*90*, *91*). We profiled three independent fragments per genotype per treatment across all three host-symbiont pairs (1:C, 2:D, 3:D; N = 18 total samples). For these samples, the V4 region of the 16S rRNA gene was amplified and sequenced following standard protocols (see Supplemental Methods).

Quality filtering, denoising, merging, and taxonomy assignments of 16S rRNA gene reads were conducted with DADA2 (*92*) against the Silva v.138.1 database (*93*) and the National Center for Biotechnology Information nucleotide database using blast+ (*94*). ASV data assigning to chloroplast, mitochondria, or non-bacterial were removed, and then contamination was removed using negative controls and the decontam package (*95*).

Alpha diversity indices (Shannon index, Simpson’s index, ASV richness, and evenness) were calculated on the full cleaned ASV data set (non-rarefied) using the estimate richness function in Phyloseq (*78*). Bray–Curtis dissimilarity-based community distance matrices were also computed using principal coordinates analysis (PCoA) with Phyloseq (*78*) and permutational multivariate analysis of variance (PERMANOVA) to analyze microbiome composition and structure differences among treatments. Taxonomic composition was characterized at class and family levels. Relative abundance differences between temperature treatments were compared using an ANOVA (aov) with treatment and genotype as fixed effect. All read tracking statistics are in table S15. The sequence reads can be found in the NCBI Short Read Archive (SRA) database as BioProject accession PRJNA1314813.

### Pathogen challenge mesocosm experiment

To test how different host-symbiont pairings performed when exposed to a pathogen, a pathogen challenge experiment using *Vibrio coralliilyticus* was performed on three genotypes (1:C, 2:D, and 3:D), with n = 12 fragments per genotype per treatment, yielding a total of N = 72 fragments. In brief, infection experiments were performed under control lighting conditions (photosynthetic photon flux density = ∼80 µmol m−2 s−1 on a 7.5:16.5h L:D cycle) and a constant temperature of 29°C in a 10 L aquarium. 29°C was used to activate the virulence of *V. coralliilyticus* without inducing coral physiological stress. To inoculate the pathogenic bacteria, fragments were transferred from their holding tanks into sterile glass jars and immediately inoculated with 15 μl of the bacterial suspension (∼10⁷ cells) without disturbing coral tissue. After one minute, treated fragments were gently placed into experimental tanks maintained at 29°C, which corresponded to pathogen treatment conditions. Control fragments were treated the same, but were given 15 μl of heat-killed *V. coralliilyticus*. Tank temperatures were maintained at 29°C for the duration of the pathogen challenge experiment using Aqualogic Digital Temperature Controllers. Coral survival and tissue loss were monitored by the same observer daily for the period of seven days post inoculation. Tissue lysis was evaluated through visual inspection and calculated as a percentage, based on the estimated area of tissue loss relative to the overall live tissue. To compare differences in mortality rates between the pathogen challenge treatments, survival v.3.5-5 and survminer v.0.4.9 packages (*96*) were used. Survival probability curves were examined by performing a log-rank test to test for significant differences among the treatment groups. To compare the percentage of tissue loss data from the final time point, we used a linear mixed model to assess the interactive effects of pathogen treatment and genotype (with random effect of tank).

### Reanalysis of gene expression data from corals hosting different algal symbiont types

To assess whether gene expression patterns between *C*- vs *D*-hosting corals observed in this study were also observed in other corals, we reanalyzed gene expression datasets from two studies: (1) adult *Pocillopora grandis* colonies hosting either *Cladocopium* (*C*) or *Durusdinium* (*D*) algal symbionts (*32*), and (2) *C-* or *D*-infected *Acropora tenuis* recruits (*33*). Raw gene counts generated in each study were reanalyzed for differential gene expression (FDR < 0.05) with DESeq2 v1.44.0 (*81*). Next, overall expression profiles were compared between *C-* and *D-*hosting corals using PCA of *vst*-normalized counts with subsequent PERMANOVA testing with the adonis2 function in the vegan package (*82*). Gene Ontology enrichment analysis was conducted using Mann-Whitney U tests (*83*) to test for functional differences in GO terms associated with *Biological Processes* between *C-* vs *D*-hosting *P. grandis*. For *A. tenuis* data, only differentially upregulated genes were used for GO enrichment analysis using the topGO package (*97*). Only GO terms with p-values <0.05 were considered significantly enriched.

For the reanalysis of lesion progression in corals exposed to stony coral tissue loss disease (SCTLD), we re-examined data from (*54*) to evaluate whether algal type affects disease susceptibility. Lesion growth rate (cm² hr⁻¹) was quantified across four stony coral species (*Colpophyllia natans*, *Montastraea cavernosa*, *Orbicella annularis*, and *Porites astreoides*), each hosting distinct algal types. Coral fragments were maintained in mesocosms that either contained a healthy *Diploria labyrinthiformis* donor (control) or an SCTLD-infected donor (disease). Lesion area was tracked over time. Differences in lesion growth rate were analyzed using an ANOVA testing for effects of algal type, treatment, and the interaction of algal type x treatment.

## ACKNOWLEDGEMENTS

We thank Katelyn Mansfield, Nicola Kriefall, Brooke Benson and previous Marine Physiology and Climate Change students for early experiments related to this research. We thank Zeba Wunderlich and Davies lab members for constructive comments on this work. We thank the Boston University (BU) Marine Program staff for help with coral maintenance, and the BU Supercomputing Center for assistance with computational analysis. We thank Justin Scace (BU) for help in experimental set-up, maintenance, and troubleshooting, and we thank Julia Hammer Mendez (BU) for coordinating the BU Marine Program. Finally, we thank three anonymous reviewers whose suggestions substantially improved this manuscript.

## FUNDING

National Science Foundation grant IOS-1937650 (TDG, SWD)

NSF Research Experience for Undergraduates grant (TDG, TA, JA, OD).

BU Marine Program Warren Mcleod Summer Award, Alistair Economakis Award (JD-A)

Republic of China (Taiwan) Ministry of Education Scholarship, and BU Marine Program Warren

McLeod Summer Award (MC)

BU Marine Program (KT) BU Work study (OJ)

BU Undergraduate Research Opportunities Program (KST)

## AUTHORS CONTRIBUTIONS

Conceptualization: SWD, TDG, JD-A

Methodology: JD-A, MC, AB, JD, AL, AS, KRT, OJ, KST, WW, TA, JA, OD

Investigation: JD-A, MC, AB, JD, AL, AS, KRT, OJ, KST, WW, TA, JA, OD, TDG

Visualization: JD-A,WW, TA, JA, OD Supervision: TDG, SWD

Writing—original draft: JD-A

Writing—review & editing: JD-A, TDG, SWD

## COMPETING INTERESTS

Authors declare that they have no competing interests.

## DATA AND MATERIALS AVAILABILITY

Authors declare that they have no competing interests.

## Notes

### Competing Interest Statement

The authors have declared no competing interest.

### Summary of Updates

This revised version of the manuscript contains new figures and reanalysis of the data set.

