## Supplementary Information for "Algae-specific Immune Modulation Influences Responses to Heat and Pathogen Challenge in a Symbiotic Coral"

**Includes:**

Supplementary Methods

Ten figures

Seven Tables

Supplementary References

### Supplementary Methods

#### Coral husbandry and fragment preparation

Colonies of five distinct genotypes of *P. acuta* acquired from the aquarium trade have been maintained at Boston University for more than nine years under common garden conditions (temperature: 26 ± 1 °C, salinity: 34-35 ppt, minimum photosynthetic photon flux density:120 μmol $m^{-2}$ $s^{-1}$ on an 8:16 hour Light:Dark cycle). Corals were fed three times a week with newly hatched brine shrimp (*Artemia salina*). These coral genotypes were determined to associate predominantly with one of two Symbiodiniaceae genera: *Cladocopium* and *Durusdinium* (formerly Clade C and D, respectively) and are referred to as *C*- and *D*-hosting throughout the manuscript. All five host-symbiont pairs (1:C, 2:D, 3:D, 4:C, 5:D) were included in baseline gene expression comparisons. Due to coral husbandry challenges, all follow-up experiments and measurements were conducted on genotypes 1:C, 2:D and 3:D. Details on specific genotype inclusion and replication are explained in each relevant methods subsection.

#### RNA isolation and sequencing from *D*- and *C*-hosting corals

Total RNA was isolated from coral tissue using the RNAqueous Total RNA Isolation Kit. RNA quality was assessed by visualizing ribosomal RNA bands on agarose gels. RNA quantification was assessed using a DS11+ spectrophotometer. Three independent libraries for host-symbiont pairings 1:C, 2:D, and 3:D (total RNA ~600 ng per sample) were submitted to the University of Texas at Austin Genomic Sequencing and Analysis Facility (GSAF) for TagSeq library preparation and sequencing (single-end 100 bp) on NovaSeq6000. For host-symbiont pairings 4C and 5D, one library per genotype was prepared in-house and sequenced on the Illumina HiSeq 2500 (single-end 50 bp) at Tufts University Core Facility.

**Coral host genotype and dominant algal symbiont assessments**

Identification of host clones was conducted as described by Rivera et al. (1). Briefly, cleaned reads were mapped to the concatenated genomes of *Pocillopora acuta* (2), and an algal symbiont database (transcriptomes of *Symbiodinium spp:* (3); *Breviolum spp:* (3); *Cladocopium spp:* (4); *Durusdinium spp:* (4)) following (5) using Bowtie2 v2.5.1 (6). Reads mapping to the host genome were then used to detect single nucleotide polymorphisms (SNPs) using ANGSD v0.935 (7). A hierarchical clustering tree (hclust) of samples based on pairwise identity by state distances calculated in ANGSD determined genetic similarity, which facilitated the identification of host clones (fig. S8a). To characterize the relative proportions of reads belonging to each algal symbiont genera in each library, reads mapping to the algal symbiont database were used to calculate the relative abundances of transcripts mapping to each algal type. All samples had 98.1 ± 0.017 % (Mean algal transcript fraction ± SD) of reads mapping to a single genus, which allowed for the identification of *Cladocopium*- and *Durusdinium*-dominated individuals (fig. S8b).

**Baseline gene expression profiling of *C*- and *D*-hosting corals**

Raw reads were processed with the TagSeq pipeline (https://github.com/z0on/tag-based_RNAseq). Briefly, Fastx_toolkit trimmed adapters and poly(A)+ tails, and sequences shorter than 20 base pairs were removed. Reads that passed quality filtering were then mapped to concatenated genomes of *Pocillopora acuta* (host genome, Stephens et al. 2022), *Durusdinium trenchii* (8), and *Cladocopium goreaui* (9) with Bowtie2 v2.5.1 (6). Reads mapping to the host genome were used in downstream analyses. Outliers were determined using the arrayQualityMetrics package (10), where an outlier was considered as a sample that failed two or more tests; no outliers were detected. To remove the effect of sequencing platform, a batch effect correction was applied on raw host count data using ComBat-seq (11), specifying sequencing platform as batch and symbiont type as the variable of interest. Differential gene expression analysis of *C*- vs *D*-hosting corals was performed using DESeq2 v1.44.0 with an FDR<0.05 (12). Expression data were *rlog*-normalized using the vegan package (13) and then a Principal Component Analysis (PCA) was conducted to characterize overall variation in gene expression among *C*- and *D*-hosting corals. A PERMANOVA tested for significant differences in clustering using the adonis2 function in vegan (13).

**Protein extraction for Western blotting of NF-κB in *C*- vs *D*-hosting corals at baseline conditions**

Coral tissues from four samples from each of two host-symbiont pairings (1:C, 3:D; N=6 total samples) were homogenized in 100 µl of 2X SDS sample buffer (0.125 M Tris-HCl, pH 6.8, 4.6% w/v SDS, 20% w/v glycerol, 10% v/v β-mercaptoethanol, and 0.2% w/v bromophenol blue) with a mortar and pestle, and then incubated at 95°C for 10 min. Homogenates were centrifuged at 13,000 rpm for 10 min, and the supernatant was collected. Protein quantification was performed using the Qubit™ Protein BR Assay Kit (Thermo Fisher) according to the manufacturer's protocol, with a slight modification in which each sample was diluted 1:20 prior to quantification.

**Analysis of holobiont physiology**

To assess algal symbiont photochemical efficiency of Photosystem II (Fv/Fm), corals were dark-acclimated for at least 30 min, then Fv/Fm was measured daily throughout the experiment using a Walz Junior-PAM™. Three replicate Fv/Fm measurements were taken from three random midbranch points for each fragment and averaged within each time point. To estimate changes in red channel intensity, a proxy for bleaching (14), photos of all fragments were taken on Day 0 and Day 12 on an iPhone 13 to assess coral color changes throughout the experiment. Photos were white-balanced in Photoshop version 25.1 and then uploaded to MATLAB for color intensity analysis following Winters et al. (14). In brief, ten random points on the coral in each photograph were selected, and red channel intensities were quantified and averaged for each coral fragment.

To estimate algal density from coral fragments at the end of the experiment, coral tissue was removed from skeletons via airbrushing using an airgun with sterile saltwater. Tissue slurry was homogenized for 3 min using a handheld tissue homogenizer (Tissue-Tearor Model 985370, BioSpec Products, Inc.), followed by centrifugation at 4,400 rpm for 3 min to separate coral host and algal components. Supernatant was collected and represents the coral host fraction for physiology measurements. Host slurries were bead blasted using Sigma glass beads and a Fisher Scientific Bead Mill 24 at 6 m/s for 2 min for other physiological measurements. The symbiont pellet was resuspended in 1 ml of sterile seawater. Symbiont densities (cells/cm2) were measured for three replicate 10 µl tissue slurry aliquots per fragment using a hemocytometer under a light microscope. Counts were averaged and normalized to live coral tissue surface area.

To quantify symbiont chlorophyll-*a* and chlorophyll-c2 pigments ($\mu g/cm^2$), symbiont tissue extracts were centrifuged, and the resulting pellets were left overnight at 4°C in 1 ml of 90% acetone in the dark. Sample absorbance at 630 and 663 nm (three 200 µl aliquots of each sample) were measured on a Synergy H1™ Microplate Reader spectrophotometer. Pigment concentrations were calculated according to the equations formulated by Jeffrey and Haxo (1968): chlorophyll-a $= 13.31(\text{Absorbance}_{663}) - 0.27(\text{Absorbance}_{630})$; chlorophyll-c2 $= -8.37(\text{Absorbance}_{663}) + 51.72(\text{Absorbance}_{630})$. To adjust for the reduced assay volume (200 µl) compared to the original volume (1 ml), a correction factor was used and the resulting values were normalized to the total original slurry volume and surface area of the living coral.

Tissue surface area calculations were conducted by 3D scanning coral skeletal fragments using an EIN-SCAN SE with the HDR and texture scanning options enabled. Each fragment was scanned three times to produce a complete watertight mesh of the coral surface. Scans were then imported into MeshLab™ software. Final photographs of skeletal areas containing live tissue were selected using the z-paintbrush tool in MeshLab to calculate the total live surface area.

Total host protein ($mg/cm^2$) was determined using a Bradford assay with Bovine Serum Albumin (BSA) standards. Each sample (6.4 µl) was diluted in artificial seawater (73.6 µl), and then equal parts (80 µl) Bradford reagent was added. Following adequate mixing, the absorbance at 595 nm was read using a Synergy H1™ Microplate Reader spectrophotometer. Protein amount was normalized to total original slurry volume and coral surface area.

#### Gene expression data processing and analysis of *C*-and *D*-hosting corals under heat stress

Quality trimming, host count generation, and outlier detection were performed following methods outlined above and all analyses described here were performed on baseline and heat-treated samples together (N=18). No outliers were detected. To remove the main effect of host genotype, which is known to be a primary driver of gene expression (15), a batch effect correction was applied on raw host count data using ComBat-seq (11), specifying genotype as batch and temperature as the variable of interest (Fig. S9A-C).

#### Weighted correlation network analysis (WGCNA)

Weighted Gene Correlation Network Analysis (WGCNA; (16)) was performed with genes with low basemean values (<3) removed, and remaining data *vst*-normalized using DESeq2 (12). Outlier samples were checked within the WGCNA package, and no outliers were detected. Signed connectivity among genes was determined, and eigengene expression of these modules was correlated to temperature treatments and physiological traits (minModuleSize=95, MEDissThres= 0.25). GO enrichment analysis of each module that was highly correlated to heated treatments was performed as described above with the modification of using Fisher's exact tests (presence/absence in a module) instead of continuous ranked p-values. We then conducted a manual search for Tumor Necrosis Factor (TNF), MAPK, and NF-κB (17), and symbiosis/stress genes (18) in modules associated with heat treatments to create a heatmap using pheatmap package in R.

**Microbiome data processing and analysis of *C*- and *D*-hosting corals under thermal challenge**

The V4 region of the 16S rRNA gene (~10 ng/ul) was amplified using Hyb515F (19) and Hyb806R (20) primers. The PCR reaction contained 1 μl of gDNA template, 0.025 U ExTaq, 1x ExTaq buffer (Takara), 1 μM primers, 0.2 mM of dNTPs, and molecular biology grade water in a total volume of 20 μl. PCR conditions were as follows: 95°C for 5 min, and 35 cycles of 1 min at 95°C, 2 min at 62°C, 2 min at 72°C, and 72°C for 10 min. Samples were purified using the GeneJet PCR Purification Kit (Thermo Fisher) and underwent a second PCR for dual-indexing (6 cycles using the same PCR conditions as above). Samples were pooled in equimolar concentration and sequenced at Tufts University Core Facility using Illumina MiSeq2500 (250 bp paired-end reads).

Demultiplexed raw reads were pre-processed with bbmap (21) to retain only sequences with 16S primers. Cutadapt (22) trimmed primers and amplicon sequence variants (ASVs) were identified and quality filtered with DADA2 (23). ASVs were then assigned taxonomy against the

Silva v.138.1 database (24) and the National Center for Biotechnology Information nucleotide database using blast+ (25). ASV data assigning to chloroplast, mitochondria, or non-bacterial were removed, and then contamination was removed using negative controls and the decontam package (26). Counts were rarefied to the nearest minimum number of sequence reads (1,500 per sample, Fig. S10) with *vegan* (13). Trimmed and rarefied ASVs were then subsequently filtered to eliminate low-abundance ASVs (<0.01%) using MCMC.OTU (27) and *vegan* (13).

**_Vibrio coralliilyticus_ culture preparation for pathogen challenge mesocosm experiment**

A culture of *Vibrio coralliilyticus*, a known cnidarian pathogen (28), was grown overnight (~16 h) at 30°C in Zobell Marine Broth from glycerol stock. The bacterial culture was then incubated at 30°C for 1 h without shaking to allow non-motile bacteria to settle. Motile bacteria were pelleted by centrifugation (3500×g, 5 min) and washed three times using 50 ml of filter-sterilized artificial seawater (FSASW). After the final wash, the bacterial pellet was resuspended in 1 ml FSASW. In brief, infection experiments were performed under control lighting conditions (photosynthetic photon flux density = ∼80 μmol m−2 s−1 on a 7.5:16.5h L:D cycle) and a constant temperature of 29°C in a 10 L aquarium. 29°C was used to activate the virulence of *V. coralliilyticus* without inducing coral physiological stress.

### Supplementary Figures and Tables

**Figure S1**

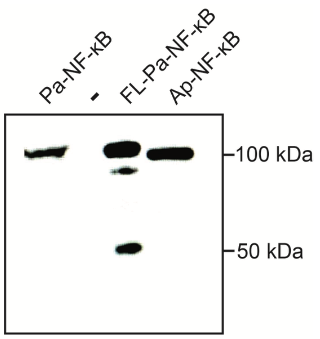


**Figure S1. Validation of anti-Aiptasia NF-κB (anti-Ap-NF-κB) cross-reactivity with *P. acuta* NF-κB (Pa-NF-κB).** Western blot using anti-Ap-NF-κB on fresh *P. acuta* tissue lysate (first lane), and lysates from 293T cells expressing FLAG-tagged Pa-NF-κB (FL-Pa-NF-κB) or 293T cells expressing Ap-NF-κB. -, 293T cells transfected with the empty vector control. The antibody specifically recognized Pa-NF-κB, detecting the full-length (~100 kDa) and processed (~50 kDa) forms.

**Figure S2**

# A

**Pa-NF-κB amino acid sequence (919 aa)**

MATNSERQIGETLTDSFVMDLLTPGYLPDISALQVPTASYQGPYMEILEQPKQRGFRFRYPCEG
PSHGGLPGQYSEKGKKSYPSVQLCNYQGPARIVVSLVTVDEPPMPHAHSLISKNSNNGVVTVQI
GPEQGMTATFPNLGIEHVTKKMVSKVLMDRYIKMQTLHTATLNALTSGDGKVFGVAGLVDQAMV
DGDRGSFDKRLAEAVAEEESQKVRAMVEEQKQSMNLNAVRLCFQAFLPDETGAFTKALPPCISN
AVYDSKAPSASNLKICRMDRNSGCVKGGDEVYLLCDKVQKDDIEVIFYETEMDTGKKTWEDRGV
FSPTDVHRQVAIVFKTPPYWNVAIEQPVKVQLELRRKSDQETSDPVEFTYQPQMFDNEQIGA**KR**
**RK**IPHFSDYL**GGGGGGGGPGMGGAGGGGGGFNFGAFGFLAPTIGFLPNYSNSGTSQSGNQGGG**
SSSQQGQSHSSGQTHASGQTHASGQPQEADLSELAWNLAEKSSAAMRDYAATGDVRYLLAVQRH
LTAVQDDNGD**TALHLAVLNARQEVVQGLLDIMASLPESFVSEY**NFLRQ**TPLHALAAITKQPRML**
**ECLLRARANARSRDRHG**N**TAVHIACMHGDAMCLKAMLNFNVTKTVLNWQNY**QGL**TPVHLAVQAG**
**SKDVLKLLNSAGANMSAQDGTSG**K**TPLHHAVEQDNLAVAGFLILEANCDVDAITLDG**N**TPLHVA**
**AASGLKGQTALLVAAGADTTVQNSDDE**IPFDLANVAEVQEILDEDEILSTDPSQDDDLASGLTG
LALGQGDMDKLDPYVRRMMAQKLDPTTGGGADWRELGRRLGLGTLENAFAIHSSPTTQVLAQYE
AADGNIKTLRQVLYDMRRGDVLEILDGHGRQDSGF**DSGLGSQS**LSASRSEELFSYDEGFSKGSS
NSSLKKGVTSLDSTRKAGNLQPIRRHQDVC

# B

**Codon optimized Pa-NF-κB cDNA sequence**

**ATG**GCCACCAACAGCGAGAGGCAGATCGGCGAGACCCTTACAGACAGCTTCGTGATGGACCTCC
TGACCCCCGGCTACCTGCCCGACATCAGCGCCCTGCAGGTGCCCACCGCCAGCTACCAGGGGCC
CTACATGGAGATCCTGGAGCAGCCCAAGCAGAGGGGCTTCAGGTTCAGGTACCCCTGCGAGGGC
CCCTCTCATGGCGGCCTGCCCGGCCAGTACAGCGAGAAGGGCAAGAAGAGCTACCCCAGCGTGC
AGCTGTGCAACTACCAGGGCCCCGCCAGAATCGTGGTTAGCCTGGTTACCGTGGACGAACCCCC
CATGCCTCACGCCCACAGCCTGATCGGCAAGAACAGCACCAACGGCGTGGTGACCGTGCAGATC
GGCCCCGAGCAGGGCATGACCGCTACCTTCCCTAACCTCGGCATCGAGCACGTGACCAAGAAGA
TGGTGGGCAAGGTGCTGATGGACAGATACATCAAGATGCAGACCCTCCACACCGCCACCCTGAA
CGCCCTGACCAGCGGCGACGGGAAGGTGTTCGGCGTGGCCGGCCTGGTGGATCAGGCCATGGTG
GACGGCGACCGGAGCAGCTTCGACAAGCACCTGGCCGAGGCCGTGGCCGAGGAGGAGAGCCAGA
AGGTGAGGGCCATGGTTGAGGAGCAGAAACAGAGCATGAACCTGAACGCCGTGAGGCTGTGCTT
CCAGGCTTTCCTGCCCGACGAGACCGGCGCCTTCACCAAGGCCCTGCACCCCTGCATCAGCAAC
GCCGTGTACGACAGCAAGGCCCCCAGCGCCAGCAACCTCAAGATCTGCAGAATGGACAGGAACT
CTGGCTGCGTGAAGGGCGGCGACGAGGTGTACCTGCTGTGTGACAAGGTGCAAAAGGACGACAT
CGAGGTGATCTTCTACGAGACCGAGATGGAGACCGGCAAGAAGACCTGGGAGGACCGGGGCGTG
TTCAGCCCCACCGACGTGCACCGGCAGGTGGCCATTGTGTTCAAGACCCCCGCCTACTGGAACG

TGGCCATCGAGCAGCCCGTGAAGGTGCAGCTGGAGCTCAGAAGAAAGAGCGACCAGGAGACCAG
CGACCCCGTGGAGTTCACCTACCAGCCCCAGATGTTCGACAAGGAGCAGATCGGCGCCAAGAGG
AGGAAGAAGATCCCCCACTTCAGCGACTACCTCGGCGGCGGAGGGGGCGGAGGCGGACCTGGCA
TGGGAGGAGCAGGAGGAGGCGGCGGAGGCTTCAACTTCGGCGCCTTCTCTCTGCTGGCCCCCAC
CATCGGATTCCTGACCAACTACACCAATAGCGGCACCAGCCAGAGCGGCAACCAGGGCGGCGGA
TCTAGCAGCCAGCAGGGCCAGAGCCACAGCAGCGGCCAGACCCACGCTAGCGGACAGCCCCAGG
AGGCCGACTTGAGCGAGCTGGCCTGGAACCTGGCCGAGAAGAGCAGCGCTGCTATGAGAGACTA
CGCCGCCACCGGCGATGTGAGGTACCTGTTGGCCGTGCAGAGGCACCTGACAGCCGTGCAGGAC
GACAACGGCGACACCGCCCTGCACCTCGCCGTGCTGAACGCCAGACAGGAGGTGGTTCAGAGCC
TGCTGGACATCATGGCCAGCCTGCCCGAAAGCTTCGTGAGCGAGTACAACTTCCTGAGACAGAC
GCCCCTGCACCTGGCCGCTATCACCAAGCAGCCCAGGATGTTGGAATGCCTGCTGAGGGCCAGG
GCCAACGCCAGGAGCAGAGACAGACACGGCAACACCGCCGTGCATATCGCCTGCATGCACGGCG
ACGCCATGTGCCTTAAAGCCATGCTGAACTTCAACGTGACCAAGACAGTGCTGAACTGGCAGAA
TTACCAGGGCCTGACCCCTGTGCATCTGGCCGTGCTGGCCGGCAGCAAGGACGTGCTGAAGCTG
CTGAACAGCGCCGGCGCCAACATGAGCGCCCAAGACGGCACCAGCGGCAAGACCCCCCTGCATC
ACGCTGTGGAGCAGGACAACCTGGCCGTGGCTGGCTTTCTGATCCTGGAGGCTAACTGCGATGT
GGACGCCATCACCCTGGACGGCAACACCCCCCTGCACGTGGCCGCTGCCTCTGGACTGAAGGGC
CAAACCGCCCTTCTGGTGGCCGCCGGAGCCGACACCACCGTGCAGAACAGCGACGACGAGATCC
CCTTCGACCTGGCCAACGTGGCCGAGGTGCAAGAGATCCTGGACGAAGACGAGATCCTGAGCAC
CGACCCCAGCCAAGACGACGACCTGGCCAGCGGCCTGACCGGCCTGGCCCTGGGACAGGGCGAT
ATGGATAAGCTGGACCCCTATGTGAGAAGGATCATGGCCCAGAAGCTAGATCCCACAACCGGAG
GGGGCGCCGACTGGAGAGAGCTGGGCAGAAGGCTGGGCTTGGGCACCCTGGAGAACGCCTTCGC
CATCCACAGCAGCCCCACCACCCAGGTGCTGGCCCAGTATGAGGCCGCCGACGGCAACATCAAG
ACACTGAGGCAGGTGCTGTACGACATGAGGAGGGGCGACGTGCTGGAGATCCTCGACGGCCATG
GCAGGCAGGACAGCGGCTTCGACAGCGGCCTGGGCAGCCAGAGCCTGAGCGCCAGCAGGTCTGA
GGAGCTGTTCAGCTACGACGAGGGCTTTAGCAAAGGCAGCAGCACCAGCAGCCTTAAGAAGGGA
GTGACCAGCCTGGACTCTACCAGGAAGGCCGGCAACCTGCAGCCTATCAGGAGACACCAGGACG
TGTGT**<u>TGA</u>**

**Figure S2. Amino acid (A) and human cell codon-optimized nucleotide (B) sequences of the Pa-NF-κB used in this study.** In (A), the predicted (nuclear localization signal) NLS is shown in **bold red font,** the glycine-rich region (GRR) is shown in **bold blue font,** the six predicted ankyrin (ANK) repeat sequences are shown in **bold green font**, and the cluster of amino acids that contains the three serine **(<u>S</u>)** residues that are potential sites of phosphorylation by IKK is shown in **bold black font.** In (B), the initiating ATG codon is in **bold font**, and the stop codon is in **<u>bold underlined font</u>.**

**Figure S3**

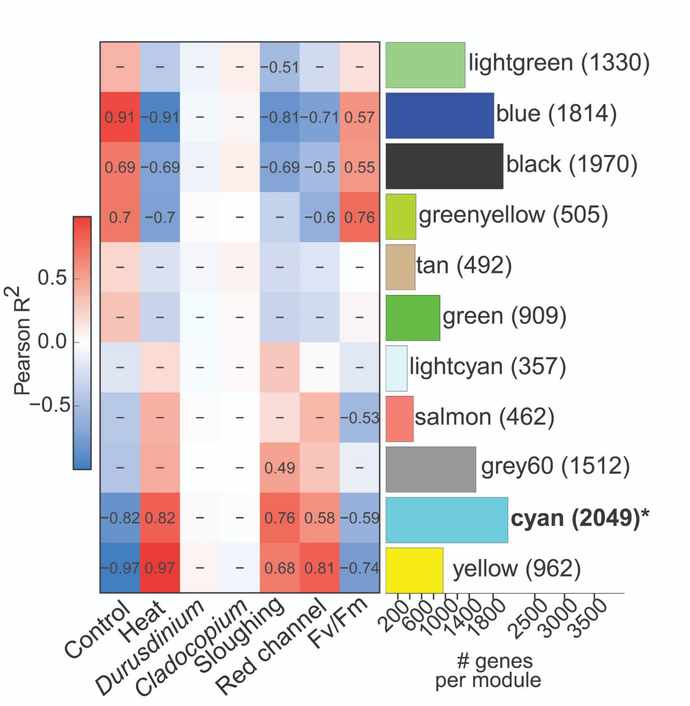


**Figure S3.** Weighted Gene Co-expression Network Analysis (WGCNA) showing Pearson correlations ($R^2$) between module eigengenes and traits, including treatment (Control, Heat), algal type (*Cladocopium*, *Durusdinium*), sloughing, Red Channel, and photosynthetic efficiency (Fv/Fm). Gene modules (denoted by colors, e.g., "lightgreen") were correlated with quantitative ("Sloughing," "Red channel," "Fv/Fm") and categorical (coded as 1 = presence, 0 = absence) traits. Each cell shows the Pearson correlation and corresponding *p*-value. Bar plots indicate the number of genes in each module.

**Figure S4**

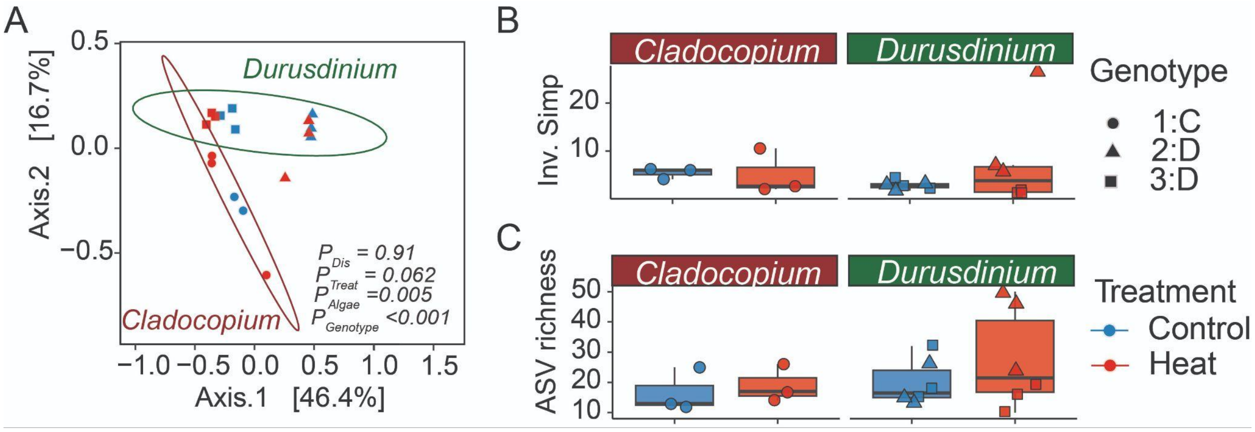


**Figure S4. Impact of heat challenge on coral microbiomes. (A)** Principal coordinates analysis based on Bray–Curtis dissimilarity of the bacterial communities (all ASVs, nonrarefied) in *C- and D*-hosting corals. Ellipses represent 95% confidence intervals. $P_{Treat}$ , $P_{Genotye}$ , $P_{Algae}$ , indicate P values for treatment, genotype, and algae. $P_{Dis}$ values indicate significant differences in dispersion between treatments. Percentages on axes indicate variation explained by each principal coordinate. (**B–C**) Mean alpha diversity indices (**B**. Inverse Simpson, **C**. ASV Richness) between control (gray) and heat (orange) treatments for *C*- and *D*-hosting corals.

**Figure S5**

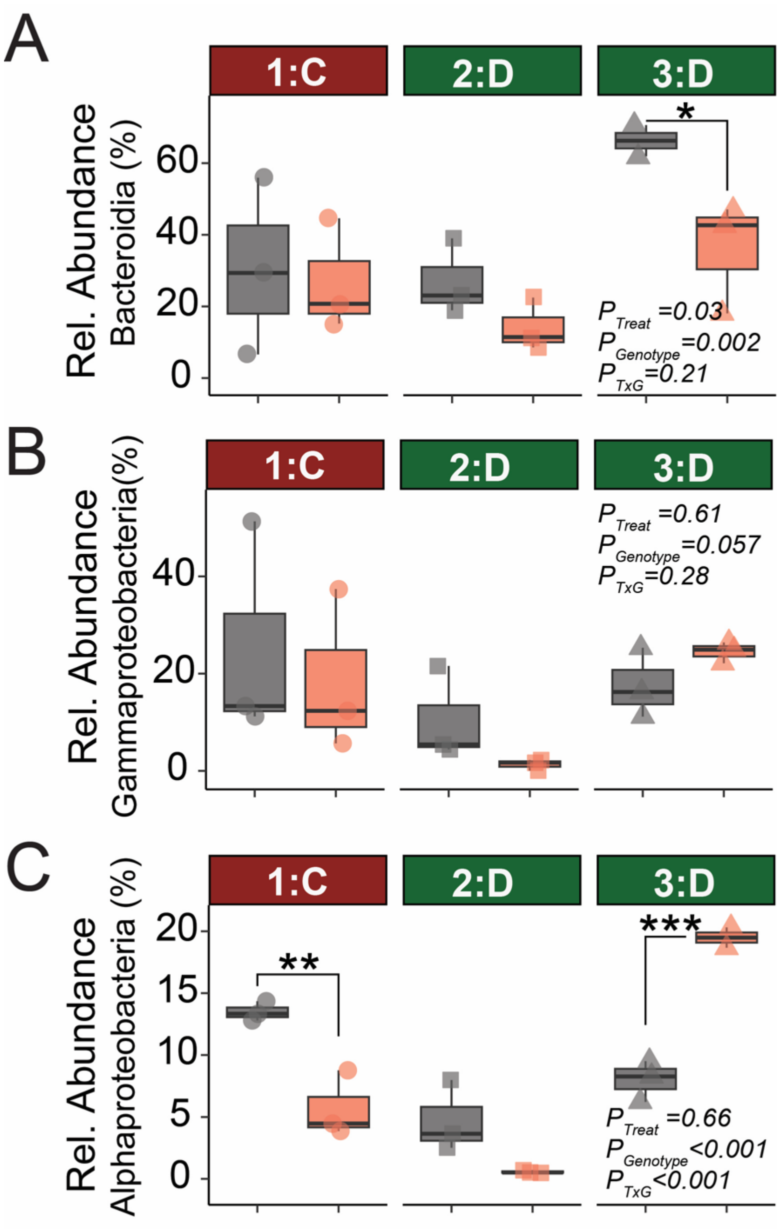


**Figure S5**. Mean relative abundances of ASVs belonging to the A) Bacteroidota, B) Gammaproteobacteria, and C) Alphaproteobacteria in *C*- and *D*-hosting corals under control (grey) and heat (orange) treatments. ANOVA results $P_{Treat}$ , $P_{Genotye}$ , $P_{TxG}$ indicate P values for treatment, genotype, and treatment x genotype.

**Figure S6**

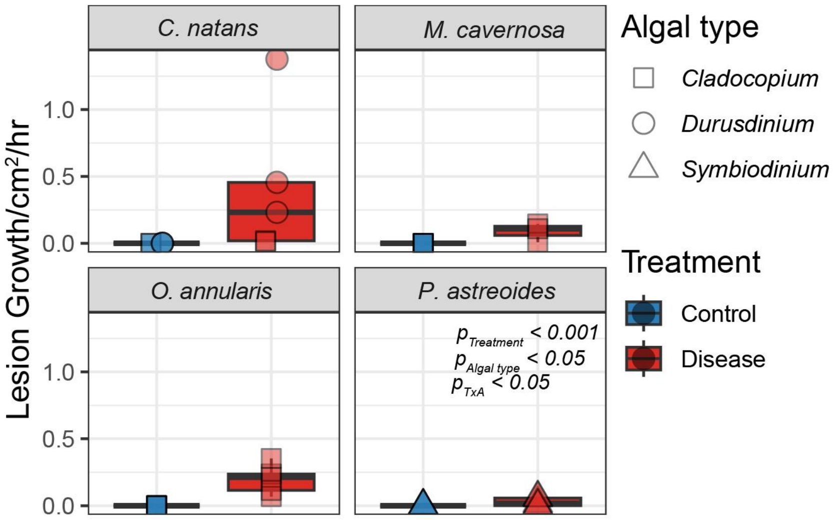


**Figure S6.** Reanalysis of lesion progression in corals exposed to stony coral tissue loss disease-infected *Diploria labyrinthiformis* (data from 29). Lesion growth rate (cm²/hr) was measured in four stony coral species, *Colpophyllia natans*, *Montastraea cavernosa*, *Orbicella annularis*, and *Porites astreoides*, hosting different algal types. Fragments were placed either in a control mesocosm with a healthy *D. labyrinthiformis* donor coral or in a disease mesocosm with an SCTLD-infected donor. *D*-hosting corals showed higher lesion growth rates.

**Figure S7**

**Mouse NF-κB p50**

PYLQILEQPKQRGFRFRYVCEGPSHGGLPGASSEKNKKSYPQVKICNYVGPAKVIVQLVTNGKN
IHLHAHSLVGKHCEDGVCTVTAGPKDMVVGFANLGILHVTKKKVFETLEARMTEACIRGYNPGL
LVHSDLAYLQAEGGGDRQLTDREKEIIRQAAVQQTKEMDLSVVRLMFTAFLPDSTGSFTRRLEP
VVSDAIYDSKAPNASNLKIVRMDRTAGCVTGGEEIYLLCDKVQKDDIQIRFYEEEENGGVWEGF
GDFSPTDVHRQFAIVFKTPKYKDVNITKPASVFVQLRRKSDLETSEPKPFLYYPEIKDKEEVQR
KRQKL

**Nv-NF-κB**

PYLEILEQPKPRGFRFRYPSEGPSHGGLPGQFSTSKSKSYPSVQVNNYQGPCRIVVTLVTKDEP
YMLHAHSLTGKNANEEGVVTVQVGPDQHMTASFPNLGIQHVTKKNVVKVLMDRFIKWQTLQNAT
FAKLSEGIKDGVDLSLFGVNTAINSNKLGFDKNVALSVANQEAAKSREYAKQQAAMDLSAVRL
CFQAYLPDQDGNFTRPLKPVYSDAVLDSKEPSASQLKICRMDKNSGCVTGGDEIYLLCDKVQKD
DIEIHFYEMDDITGKYTWEDLGKFSPCDVHRQFAIVFKTPPYWNIAIERPANVLVELRRKKKNG
GETSEPVQFTYQPQLFDKEAIGAKRRKT

**Pa-NF-κB**

PYMEILEQPKQRGFRFRYPCEGPSHGGLPGQYSEKGKKSYPSVQLCNYQGPARIVVSLVTVDEP
PMPHAHSLISKNSNNGVVTVQIGPEQGMTATFPNLGIEHVTKKMVSKVLMDRYIKMQTLHTATL
NALTSGDGKVFGVAGLVDQAMVDGDRGSFDKRLAEAVAEEESQKVRAMVEEQKQSMNLNAVRLC
FQAFLPDETGAFTKALPPCISNAVYDSKAPSASNLKICRMDRNSGCVKGGDEVYLLCDKVQKDD
IEVIFYETEMDTGKKTWEDRGVFSPTDVHRQVAIVFKTPPYWNVAIEQPVKVQLELRRKSDQET
SDPVEFTYQPQMFDNEQIGAKRRKI

**Figure S7. Amino acid sequences from the Rel Homology Domains (RHDs) used for generation of the Alphafold3 structures on DNA.** Shown are the analogous RHD amino acid sequences from the indicated proteins that were used to generate the Alphafold3 structures on the palindromic κB site (5'GGGAATTCCC3'), as shown in Fig. 2B. See ref 28) for more details on generating these structures and comparing them in PyMOL (using Root Mean Square Deviation [RMSD] values].

**Figure S8**

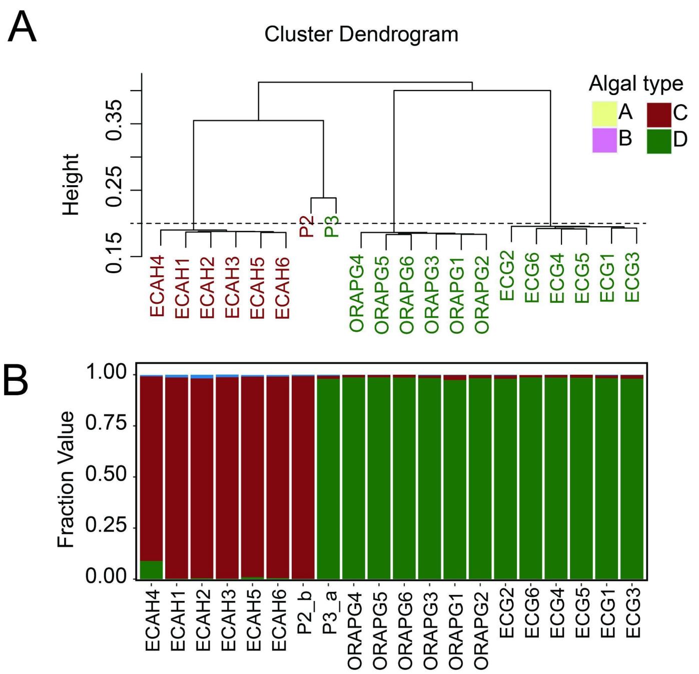


**Figure S8. Characterization of host and symbiont identity.** (**A**) Identity-by-state (IBS) dendrogram of coral fragments showing five putative genotypes. The dashed line (height = 0.20) indicates the threshold for clone assignment. Sample IDs are formatted as [Genotype][Fragment Number]. (**B**) Proportion of TagSeq reads mapping to transcriptomes of four Symbiodiniaceae genera: *Symbiodinium* (**A**), *Breviolum* (**B**), *Cladocopium* (**C**), and *Durusdinium* (**D**). Reference transcriptomes: (3) for A and B; (4) for C and D.

**Figure S9**

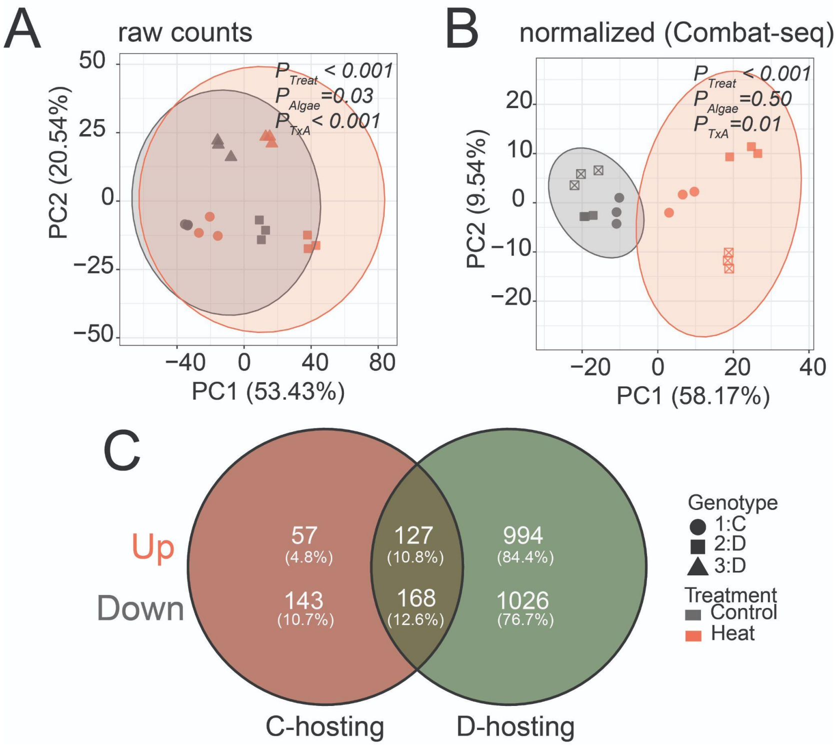


**Figure S9. Transcriptome profiling revealed host gene expression changes under heat stress.** Principal Component Analysis (PCA) of *Pocillopora acuta* based on rlog-normalized host gene expression from (**A**) raw and (**B**) ComBat-seq–corrected counts with host genotype modelled out. Ellipses represent 95% confidence intervals for each treatment group. P-values indicate significant differences from PERMANOVA; only significant terms are shown (Treat = treatment, Algae = algal type, TxA = treatment × algal type). (**C**) Venn diagram of differentially expressed host genes (DEGs; FDR < 0.05) in heated vs. control *P. acuta*. Shapes denote genotypes. *D*-hosting corals exhibited significantly more DEGs than *C*-hosting corals under heat stress (FDR < 0.05).

**Figure S10**

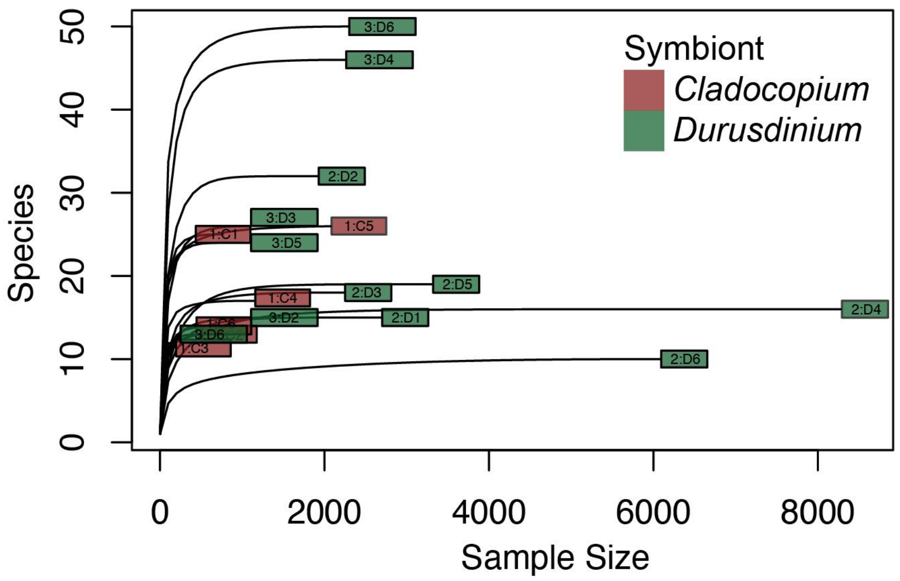


**Figure S10.** Rarefaction curves showing bacterial species richness as sequencing depth increases for *C*-hosting (brown) and *D*-hosting (green) corals under control and heat treatments. The curves plateau in both groups, indicating sufficient sequencing depth to capture most bacterial diversity in *P. acuta* hosting either *Cladocopium* or *Durusdinium*.

**Table S1**

**Table S1.** Summary of differential gene expression analysis for *C*- and *D*-hosting corals under control and heat treatments. The table shows differentially expressed genes (*FDR* < 0.05) associated with the Tumor Necrosis Factor (TNF) pathway. DE Pairwise comparison of heated vs control using raw or batch-corrected (CombatSeq) counts were shown.

| | | | | Raw data | | | | CombatSeq | | | |
|---|---|---|---|---|---|---|---|---|---|---|---|
| | | | | *D*-hosting | | *C*-hosting | | *D*-hosting | | *C*-hosting | |
| TranscriptID (Pocillopora_acuta_Hlv2___) | Gene | Description | Pathway | HeatvsCont (logFC) | HeatvsCont (padj) | HeatvsCont (logFC) | HeatvsCont (padj) | HeatvsCont (logFC) | HeatvsCont (padj) | HeatvsCont (logFC) | HeatvsCont (padj) |
| RNAseq.3335_t | TRAF3 | thioesterase binding | TNF | 0.53058132 | 0.60447515 | -0.0014971 | 0.99930992 | 0.33281181 | 0.8607913 | -0.087544 | 0.97981368 |
| RNAseq.g15629.t1 | TRAF3 | thioesterase binding | TNF | 0.56801584 | 0.37741205 | 0.43752793 | 0.76674186 | 0.60106608 | 0.43061151 | 0.32122381 | 0.85479383 |
| RNAseq.g22746.t1 | TRAF3 | thioesterase binding | TNF | 1.17548128 | 0.00846837 | 1.27926674 | 0.08937405 | 1.19219449 | 0.02957741 | 1.16777621 | 0.05952044 |
| RNAseq.g24949.t1 | TRAF3 | thioesterase binding | TNF | -0.6632232 | 0.35751977 | -0.1246357 | 0.95506211 | -0.5941062 | 0.46340293 | -0.1231871 | 0.96039909 |
| RNAseq.g25491.t1 | TRAF3 | thioesterase binding | TNF | 0.18199362 | 0.86138615 | 0.24349389 | 0.92439622 | 0.0727471 | 0.96414044 | 0.27153256 | 0.90728946 |
| RNAseq.g25494.t1 | TRAF3 | thioesterase binding | TNF | 0.78210624 | 0.18419839 | 0.19778994 | 0.93907538 | 0.81150003 | 0.30822615 | 0.15127052 | 0.94751873 |
| RNAseq.3335_t | TRAF3 | thioesterase binding | TNF | 0.53058132 | 0.60447515 | -0.0014971 | 0.99930992 | 0.33281181 | 0.8607913 | -0.087544 | 0.97981368 |
| TS.g31.t1 | MAP3K7 | MAP kinase kinase kinase activity | TNF | 0.16153317 | 0.86714999 | -0.4243328 | 0.85164867 | 0.18727347 | 0.88985928 | -0.4445216 | 0.8111462 |
| TS.g8240.t3 | AP1G1 | positive regulation of natural killer cell degranulation | TNF | 0.04604049 | 0.89143488 | -0.1767175 | 0.80515021 | 0.02831226 | 0.9501366 | -0.1868562 | 0.75751385 |
| TS.g13419.t1 | JUN | cellular response to potassium ion starvation | TNF | 1.3623862 | 0.03746047 | 0.76357476 | 0.53760393 | 1.05715011 | 0.19056742 | 0.87259937 | 0.44242449 |
| TS.g15122.t3 | MAP2K7 | JUN kinase kinase activity | TNF | -0.0829502 | 0.90281407 | -0.0024106 | 0.99906982 | -0.1168022 | 0.88360954 | 0.13051169 | 0.93329268 |
| RNAseq.g21494.t1 | DNM1L | BH2 domain binding | TNF | 0.70748865 | 0.13723352 | 0.18962534 | 0.90610301 | 0.68719863 | 0.19441579 | 0.15177532 | 0.9290624 |
| RNAseq.g13939.t1 | RPS6KA5 | ribosomal protein S6 kinase | TNF | 0.32860146 | 0.56271149 | -0.3962306 | 0.77226438 | 0.30805627 | 0.68538803 | -0.3949265 | 0.73946252 |
| RNAseq.g17589.t1 | BCL3 | B-cell CLL lymphoma 3 | TNF | 0.82452448 | 0.01190204 | 0.76046418 | 0.09312061 | 0.80629786 | 0.01996935 | 0.72374321 | 0.12771984 |
| RNAseq.g17592.t1 | BCL3 | B-cell lymphoma 3 protein | TNF | -0.4797985 | 0.56218288 | 0.0398931 | 0.98439162 | -0.3889503 | 0.65898272 | 0.14210016 | 0.94540832 |
| RNAseq.g26123.t1 | CASP7 | BIR domain binding | TNF | 0.68002574 | 0.11925532 | -0.1669696 | 0.93503092 | 0.62342306 | 0.25781087 | -0.1485233 | 0.93386183 |
| RNAseq.g9096.t1 | IFIH1 | C-terminal domain of RIG-I | TNF | 0.79601133 | 0.00222564 | 0.44214454 | 0.37724173 | 0.73695888 | 0.00429019 | 0.41820824 | 0.34184192 |

**Table S2**

**Table S2.** Summary of differential gene expression analysis for *C*- and *D*-hosting corals under control and heat treatments. The table shows differentially expressed genes (*FDR* < 0.05) associated with the Mitogen-Activated Protein Kinase (MAPK) Pathway. DE Pairwise comparison of heated vs control using raw or batch-corrected (CombatSeq) counts were shown.

| | | | | Raw data | | | | CombatSeq | | | |
|---|---|---|---|---|---|---|---|---|---|---|---|
| | | | | *D*-hosting | | *C*-hosting | | *D*-hosting | | *C*-hosting | |
| **TranscriptID** (Pocillopora_acuta_HIv2___) | Gene | Description | Pathway | HeatvsCont (logFC) | HeatvsCont (padj) | HeatvsCont (logFC) | HeatvsCont (padj) | HeatvsCont (logFC) | HeatvsCont (padj) | HeatvsCont (logFC) | HeatvsCont (padj) |
| RNAseq.g25813.t1 | NFKB1 | Nuclear factor NF-kappa-B p105 subunit | MAPK | 1.264357 | 0.00245892 | 0.68076601 | 0.41709584 | 1.2134145 | 0.01241093 | 0.63677543 | 0.43592491 |
| RNAseq.g1648.t1 | TRAF6 | tumor necrosis factor receptor binding | MAPK | 0.60398588 | 0.01102743 | 0.98491298 | 0.00042394 | 0.63068501 | 0.00490528 | 0.63677543 | 0.43592491 |
| RNAseq.g7626.t1 | MYD88_TLR | MyD88-dependent toll-like receptor signaling pathway | MAPK | -1.9318955 | 0.02543507 | 0.33403917 | 0.90402667 | -1.7196639 | 0.05722958 | 0.69576678 | 0.01752091 |
| TS.g14245.t1 | MYD88_TLR | MyD88-dependent toll-like receptor signaling pathway | MAPK | -0.1236805 | 0.8611786 | -0.4406543 | 0.72738602 | 0.01245056 | 0.98846573 | 0.41844235 | 0.86645669 |
| TS.g8240.t3 | AP1G1 | positive regulation of natural killer cell degranulation | MAPK | 0.04604049 | 0.89143488 | -0.1767175 | 0.80515021 | 0.02831226 | 0.9501366 | -0.4738311 | 0.61316306 |
| TS.g13419.t1 | JUN | cellular response to potassium ion starvation | MAPK | 1.3623862 | 0.03746047 | 0.76357476 | 0.53760393 | 1.05715011 | 0.19056742 | -0.1868562 | 0.75751385 |
| TS.g15122.t3 | MAP2K7 | JUN kinase kinase activity | MAPK | -0.0829502 | 0.90281407 | -0.0024106 | 0.99906982 | -0.1168022 | 0.88360954 | 0.87259937 | 0.44242449 |
| RNAseq.g17423.t2 | MAP4K4 | MAP kinase kinase kinase kinase activity | MAPK | 1.11082894 | 0.0992134 | -0.2570484 | 0.92439622 | 0.96974525 | 0.27222191 | 0.13051169 | 0.93329268 |
| RNAseq.g26359.t1 | MYC | V-myc avian myelocytomatosis viral oncogene homolog | MAPK | -0.1468017 | 0.79717943 | 0.51338508 | 0.35726482 | 0.0187499 | 0.98144779 | -0.2669489 | 0.91359664 |
| RNAseq.g26769.t1 | MYCBPAP | MYCBP associated protein | MAPK | -0.0087189 | 0.9848409 | 0.02434839 | 0.98638153 | -0.0214591 | 0.97134295 | 0.57824036 | 0.25455752 |
| RNAseq.g7674.t1 | GRB2 | neurotrophin TRKA receptor binding | MAPK | 0.54095539 | 0.01146679 | -0.0429959 | 0.97155683 | 0.54877155 | 0.01167515 | 0.00207232 | 0.99930059 |
| RNAseq.g2232.t1 | FGFR2 | Fibroblast growth factor receptor 2 | MAPK | 1.93705941 | 00001858959 | 0.69942457 | 0.50778574 | 1.86523911 | 0.00019758 | -0.0623405 | 0.95620836 |
| RNAseq.g27408.t1 | FGF7 | Fibroblast growth factor 7 | MAPK | 1.46577368 | 0.00594615 | -0.5054337 | 0.83548044 | 1.49495041 | 0.05207219 | 0.68758942 | 0.3617778 |
| RNAseq.g3781.t1b | FGF20 | fibroblast growth factor | MAPK | -0.2412988 | 0.50034236 | -0.2504471 | 0.78784605 | -0.2732123 | 0.47449656 | -0.5077216 | 0.77803701 |
| TS.g28704.t1 | FGF18 | positive regulation of cell chemotaxis to fibroblast | MAPK | -1.52869 | 0.00335814 | -0.7139278 | 0.26596943 | -1.348208 | 0.00582088 | -0.2735262 | 0.7213205 |
| TS.g27984.t1 | MAP3K13 | MAP kinase kinase kinase activity | MAPK | -0.4352887 | 0.77133611 | 0.1089746 | 0.96245363 | -0.4526894 | 0.78353386 | -0.6830428 | 0.37557194 |
| RNAseq.g14886.t1 | MAP3K2 | Mitogen-activated protein kinase kinase kinase | MAPK | 0.56735476 | 0.46456754 | 0.37582902 | 0.82188345 | 0.46306129 | 0.66472992 | 0.36628066 | 0.87976496 |
| RNAseq.g15261.t1 | MAP3K2 | Mitogen-activated protein kinase kinase kinase | MAPK | 0.49858617 | 0.40604172 | 0.19921833 | 0.92009871 | 0.51016775 | 0.45252101 | 0.34149857 | 0.87413503 |
| RNAseq.g15322.t1 | MAP3K2 | MAP kinase kinase kinase activity | MAPK | -0.3836216 | 0.60606965 | 0.49695397 | 0.7296952 | -0.3867773 | 0.67478801 | 0.16378121 | 0.93191167 |
| RNAseq.g363.t1 | MAP3K20 | MAP kinase kinase kinase activity | MAPK | 0.33692231 | 0.47333537 | 0.50594747 | 0.54642729 | 0.34666988 | 0.54728732 | 0.47164404 | 0.74273805 |
| RNAseq.g15261.t1 | MAP3K2 | Mitogen-activated protein kinase kinase kinase | MAPK | 0.49858617 | 0.40604172 | 0.19921833 | 0.92009871 | 0.51016775 | 0.45252101 | 0.46637229 | 0.52827532 |
| TS.g22447.t2 | MAPK7 | negative regulation of ERK5 cascade | MAPK | 1.88007691 | 0.00078238 | 0.91728817 | 0.47017965 | 1.78297019 | 0.012856 | 0.16378121 | 0.93191167 |
| RNAseq.g24760.t1 | PLA2G4A | Phospholipase A2 | MAPK | -0.697456 | 0.42168484 | 0.09169335 | 0.97278272 | -0.6462624 | 0.48639578 | 0.85077706 | 0.46642655 |
| RNAseq.g26356.t1 | PLA2G4A | Phospholipase A2 | MAPK | 2.60271703 | 0 | 1.43818938 | 0.0001201 | 2.71709517 | 3.55E-26 | 0.15256537 | 0.9554846 |

**Table S3**

**Table S3.** Summary of differential gene expression analysis for *C*- and *D*-hosting corals under control and heat treatments. The table shows differentially expressed genes (*FDR* < 0.05) associated with the nuclear factor-kappa B (NF-κB) Pathway. DE Pairwise comparison of heated vs control using raw or batch-corrected (CombatSeq) counts were shown.

| TranscriptID (Pocillopora_acuta_HIv2___) | Gene | Description | Pathway | Raw data | | | | CombatSeq | | | |
|---|---|---|---|---|---|---|---|---|---|---|---|
| | | | | *D*-hosting | | *C*-hosting | | *D*-hosting | | *C*-hosting | |
| | | | | HeatvsCont (logFC) | HeatvsCont (padj) | HeatvsCont (logFC) | HeatvsCont (padj) | HeatvsCont (logFC) | HeatvsCont (padj) | HeatvsCont (logFC) | HeatvsCont (padj) |
| RNAseq.g25813.t1 | NFKB1 | Nuclear factor NF-kappa-B p105 subunit | NFKB | 1.264357 | 0.00245892 | 0.68076601 | 0.41709584 | 1.2134145 | 0.01241093 | 0.63677543 | 0.43592491 |
| RNAseq.3335_t | TRAF3 | thioesterase binding | NFKB | 0.53058132 | 0.60447515 | -0.0014971 | 0.99930992 | 0.33281181 | 0.8607913 | 0.63677543 | 0.43592491 |
| RNAseq.g14334.t1 | TRAF3 | TNF receptor-associated factor | NFKB | 1.13536686 | 0.23541489 | #N/A | #N/A | 1.09157198 | 0.43852407 | -0.087544 | 0.97981368 |
| RNAseq.g15629.t1 | TRAF3 | thioesterase binding | NFKB | 0.56801584 | 0.37741205 | 0.43752793 | 0.76674186 | 0.60106608 | 0.43061151 | #N/A | #N/A |
| RNAseq.g22746.t1 | TRAF3 | thioesterase binding | NFKB | 1.17548128 | 0.00846837 | 1.27926674 | 0.08937405 | 1.19219449 | 0.02957741 | 0.32122381 | 0.85479383 |
| RNAseq.g24949.t1 | TRAF3 | thioesterase binding | NFKB | -0.6632232 | 0.35751977 | -0.1246357 | 0.95506211 | -0.5941062 | 0.46340293 | 1.16777621 | 0.05952044 |
| RNAseq.g25491.t1 | TRAF3 | thioesterase binding | NFKB | 0.18199362 | 0.86138615 | 0.24349389 | 0.92439622 | 0.0727471 | 0.96414044 | -0.1231871 | 0.96039909 |
| RNAseq.g25494.t1 | TRAF3 | thioesterase binding | NFKB | 0.78210624 | 0.18419839 | 0.19778994 | 0.93907538 | 0.81150003 | 0.30822615 | 0.27153256 | 0.90728946 |
| RNAseq.3335_t | TRAF3 | thioesterase binding | NFKB | 0.53058132 | 0.60447515 | -0.0014971 | 0.99930992 | 0.33281181 | 0.8607913 | 0.15127052 | 0.94751873 |
| TS.g31.t1 | MAP3K7 | MAP kinase kinase kinase activity | NFKB | 0.16153317 | 0.86714999 | -0.4243328 | 0.85164867 | 0.18727347 | 0.88985928 | -0.087544 | 0.97981368 |
| RNAseq.g1616.t1 | TRAF6 | TNF receptor-associated factor 6 | NFKB | 0.39718714 | 0.49367929 | 0.37381288 | 0.77507569 | 0.38512646 | 0.6045179 | -0.4445216 | 0.8111462 |
| RNAseq.g1619.t1 | TRAF6 | TNF receptor-associated factor 6 | NFKB | 0.96163188 | 0.57414993 | 0.5528299 | 0.47507577 | 0.49353096 | 0.8490075 | 0.23238218 | 0.88670786 |
| RNAseq.g1648.t1 | TRAF6 | tumor necrosis factor receptor binding | NFKB | 0.60398588 | 0.01102743 | 0.98491298 | 0.00042394 | 0.63068501 | 0.00490528 | 1.03539787 | 0.41137359 |
| RNAseq.g7626.t1 | MYD88_TLR | MyD88-dependent toll-like receptor signaling pathway | NFKB | -1.9318955 | 0.02543507 | 0.33403917 | 0.90402667 | -1.7196639 | 0.05722958 | 0.69576678 | 0.01752091 |
| TS.g14245.t1 | MYD88_TLR | MyD88-dependent toll-like receptor signaling pathway | NFKB | -0.1236805 | 0.8611786 | -0.4406543 | 0.72738602 | 0.01245056 | 0.98846573 | 0.41844235 | 0.86645669 |
| RNAseq.g9482.t1 | DDX4 | Belongs to the DEAD box helicase family | NFKB | -1.4713763 | 0.0406948 | 0.28059014 | 0.89158556 | -1.2923889 | 0.09872486 | -0.4738311 | 0.61316306 |
| RNAseq.g8804.t1 | MALT1 | lymphoid tissue lymphoma translocation | NFKB | -0.2663331 | 0.61709085 | -0.3206789 | 0.73948922 | -0.2979466 | 0.61614217 | 0.34786224 | 0.87549898 |

### Table S4

**Table S4.** Summary of differential gene expression analysis for *C*- and *D*-hosting corals under control and heat treatments. The table shows differentially expressed genes (*FDR* < 0.05) associated with genes associated with stress. DE Pairwise comparison of heated vs control using raw or batch-corrected (CombatSeq) counts were shown.

| TranscriptID (Pocillopora_acuta_Hlv2___) | Gene | Description | Raw data | | | | CombatSeq | | | |
|---|---|---|---|---|---|---|---|---|---|---|
| | | | *D*-hosting | | *C*-hosting | | *D*-hosting | | *C*-hosting | |
| | | | HeatvsCont (logFC) | HeatvsCont (padj) | HeatvsCont (logFC) | HeatvsCont (padj) | HeatvsCont (logFC) | HeatvsCont (padj) | HeatvsCont (logFC) | HeatvsCont (padj) |
| RNAseq.1271_t | LMAN1 | Lectin, mannose-binding 1 | 0.1897669234 | 0.8075551437 | 0.3556477966 | 0.7878460502 | 0.1171529 | 0.90993635 | 0.37546707 | 0.76371428 |
| RNAseq.g11210.t1 | CAT | Occurs in almost all aerobically respiring organis | 0.4388529115 | 0.3607615427 | 0.620482178 | 0.3300620443 | 0.33498525 | 0.58936525 | 0.69945415 | 0.22195276 |
| RNAseq.g14334.t1 | TRAF3 | TNF receptor-associated factor | 1.135366864 | 0.2354148949 | #N/A | #N/A | 1.09157198 | 0.43852407 | #N/A | #N/A |
| RNAseq.g16342.t1 | RCHY1 | ring finger and CHY zinc finger | #N/A | #N/A | -0.5904553125 | 0.7111558117 | #N/A | #N/A | -0.6146705 | 0.6871407 |
| RNAseq.g17086.t1 | peroxidase | peroxidase activity | 2.234299357 | 8.51968069974128e-13 | 0.3959514648 | 0.819314201 | 2.16258718 | 1.88E-08 | 0.76153305 | 0.12299832 |
| RNAseq.g17168.t1 | TRAF4 | activation of NF-kappaB-inducing kinase activity | 1.483472257 | 3.99080199163414e-06 | 0.390183109 | 0.5880572418 | 1.1467214 | 0.00011798 | 0.30739553 | 0.70509426 |
| RNAseq.g17589.t1 | BCL3 | B-cell CLL lymphoma 3 | 0.8245244838 | 0.01190204427 | 0.7604641819 | 0.09312061377 | 0.80629786 | 0.01996935 | 0.72374321 | 0.12771984 |
| RNAseq.g20357.t1 | MIB2 | E3 ubiquitin-protein ligase | 1.053453957 | 0.006123103234 | 0.7899124878 | 0.036937493 | 1.08429672 | 0.00260314 | 0.7197715 | 0.08135669 |
| RNAseq.g20359.t1 | MIB2 | E3 ubiquitin-protein ligase | 0.1764165102 | 0.6384804152 | 0.4276558908 | 0.4815624582 | 0.08426769 | 0.89187971 | 0.33695808 | 0.58685973 |
| RNAseq.g20360.t1 | MIB2 | ubiquitin-protein transferase activity | #N/A | #N/A | 1.287359843 | 0.001331231435 | 1.94159864 | 4.46E-06 | 1.26366385 | 0.00164089 |
| RNAseq.g20545.t1 | AGO2 | Required for RNA-mediated gene silencing (RN/ | 1.123469312 | 3.36705984863699e-05 | 0.3872755639 | 0.532208837 | 1.05658651 | 7.80E-05 | 0.3453378 | 0.57631169 |
| RNAseq.g20667.t1 | athione S-transfe | Glutathione S-transferase, C-terminal domain | 0.7998492845 | 0.01696043072 | 1.58894778 | 2.09214603681982e-05 | 0.72796359 | 0.04346597 | 1.56058662 | 2.38E-06 |
| RNAseq.g20668.t1 | athione S-transfe | Glutathione S-transferase, C-terminal domain | 1.656887235 | 2.74280535750831e-06 | 2.024418841 | 4.43680290303552e-06 | 1.59268759 | 2.77E-05 | 2.03483913 | 5.52E-08 |
| RNAseq.g20669.t1 | athione S-transfe | Glutathione S-transferase, C-terminal domain | 0.7795195341 | 0.0006597632409 | 0.8838806973 | 0.0004806392948 | 0.75014345 | 0.00013524 | 0.96168823 | 0.0001007 |
| RNAseq.g22605.t1 | NCAM2 | cell adhesion | -1.319149215 | 0.0116224343 | -0.09006398922 | 0.960317309 | -1.2348315 | 0.01151281 | -0.1067092 | 0.95774585 |
| RNAseq.g25735.t1 | utathione peroxida | glutathione peroxidase activity | -2.080347594 | 0.007479412239 | -0.6298125992 | 0.6968346459 | -1.9644172 | 0.01513713 | -0.6044424 | 0.70831388 |
| RNAseq.g25813.t1 | NFKB1 | Nuclear factor NF-kappa-B p105 subunit | 1.264356999 | 0.002458916228 | 0.6807660099 | 0.417095844 | 1.2134145 | 0.01241093 | 0.63677543 | 0.43592491 |
| RNAseq.g28985.t1 | CYP46A1 | Cytochrome P450, family 46, subfamily A, polyp | 1.018726496 | 0.08279020914 | 0.5133105368 | 0.8037759855 | 1.01323569 | 0.164325 | 0.49455895 | 0.75519194 |
| RNAseq.g30855.t1 | CYP21A2 | Cytochrome P450, family 21, subfamily A, polyp | 2.568557674 | 2.92125891595792e-07 | 1.12575146 | 0.5058091293 | 2.66287635 | 0.00212894 | 1.10182876 | 0.3496085 |
| RNAseq.g30858.t1 | CYP21A2 | Cytochrome P450, family 21, subfamily A, polyp | 2.178994791 | 1.83032913067762e-13 | 0.6896223717 | 0.4246838739 | 2.128055 | 1.69E-10 | 0.81741099 | 0.07689748 |
| RNAseq.g30954.t1 | peroxidase | peroxidase activity | 1.906428453 | 0.005792239521 | 0.4606052962 | 0.8279730989 | 2.01820136 | 0.07589766 | #N/A | #N/A |
| RNAseq.g6656.t1 | HSP90B1 | heat shock protein 90kDa beta (Grp94), membe | 0.6165039628 | 0.00752280138 | 0.6810804136 | 0.01276795504 | 0.65046527 | 0.00086905 | 0.66453164 | 0.01797342 |
| RNAseq.g6962.t1 | HSPA5 | Belongs to the heat shock protein 70 family | 1.08756983 | 2.21521794115082e-06 | 0.712852818 | 0.018336655 | 0.99496161 | 3.68E-06 | 0.69336928 | 0.01866228 |
| RNAseq.g7990.t1 | HSF1 | cellular response to nitroglycerin | -0.2985083305 | 0.6172564482 | 0.3818889378 | 0.7128624402 | -0.2335478 | 0.77188425 | 0.35015521 | 0.72431785 |
| RNAseq.g8399.t1 | HSPA8 | Heat shock cognate 71 kDa | 0.6470051614 | 0.0114424104 | 0.1483565411 | 0.8715803686 | 0.73432879 | 0.00118924 | 0.20425143 | 0.76812929 |
| RNAseq.g8806.t1 | Caspase domain | Caspase domain | 1.009036869 | 0.02273456768 | 0.2553389606 | 0.9104079001 | 1.06857305 | 0.08242661 | 0.28834762 | 0.87412985 |
| RNAseq.g9129.t1 | UBR4 | E3 ubiquitin-protein ligase | 0.5631115453 | 0.01154718873 | 0.5067431293 | 0.1532863086 | 0.5616323 | 0.01293399 | 0.4259838 | 0.23866665 |
| RNAseq.g9547.t1 | HSPA8 | Heat shock cognate 71 kDa protein | 0.3822132072 | 0.2614965235 | 0.4012698842 | 0.5285705783 | 0.38715244 | 0.27905139 | 0.3667615 | 0.570757 |
| TS.g11372.t1b | Gal_Lectin | Galactose binding lectin domain | -3.647210053 | 5.04E-09 | -1.966458606 | 1.58232739079214e-18 | -3.2507455 | 2.68E-10 | -1.4126531 | 0.04307199 |
| TS.g13497.t1c | HSPA4L | heat shock | 0.7112106907 | 0.0001165112475 | 0.3435820188 | 0.3564400838 | 0.68087389 | 1.21E-05 | 0.27131427 | 0.5025376 |
| TS.g14259.t1 | TRAF2 | Tnf receptor-associated factor 2 | 2.036075235 | 5.68015720389e-05 | 0.8847959938 | 0.4758786251 | 1.84766661 | 0.00147495 | 0.85274981 | 0.34802655 |
| TS.g14265.t2 | TRAF3 | Tnf receptor-associated factor 2 | 1.090196292 | 1.47213316902451e-06 | -0.07816399852 | 0.9624536349 | 1.06494703 | 1.60E-05 | 0.0361487 | 0.97921847 |
| TS.g21596.t1 | TRAF4 | activation of NF-kappaB-inducing kinase activity | -0.150249598 | 0.8121395124 | 0.6621894647 | 0.2836745599 | -0.2654559 | 0.71224715 | 0.64225451 | 0.35866228 |
| TS.g3084.t1 | Gal_Lectin | Galactose binding lectin domain | -1.543921123 | 8.0011579674301e-11 | -0.7002413935 | 0.02641608884 | -1.3995897 | 1.71E-10 | -0.669189 | 0.03081498 |
| TS.g3085.t1a | Gal_Lectin | Galactose binding lectin domain | 1.136319104 | 1.22717481742821e-08 | -0.1136067305 | 0.9726322102 | 1.08813779 | 0.00164377 | 0.28156065 | 0.75519194 |

### Table S5

**Table S5.** Summary of differential gene expression analysis for *C*- and *D*-hosting corals under control and heat treatments. The table shows differentially expressed genes (*FDR* < 0.05) associated with genes associated with symbiosis. DE Pairwise comparison of heated vs control using raw or batch-corrected (CombatSeq) counts were shown.

| | | | Raw counts | | | | CombatSeq | | | |
|---|---|---|---|---|---|---|---|---|---|---|
| | | | *D*-hosting | | *C*-hosting | | *D*-hosting | | *C*-hosting | |
| **TranscriptID** (Pocillopora_acuta_Hlv2___) | **Gene** | **Description** | HeatvsCont (logFC) | HeatvsCont (padj) | HeatvsCont (logFC) | HeatvsCont (padj) | HeatvsCont (logFC) | HeatvsCont (padj) | HeatvsCont (logFC) | HeatvsCont (padj) |
| RNAseq.g14257.t1 | NPC2 | intracellular cholesterol transport | -1.898723943 | 0.1776692645 | -0.1546808935 | 0.9456576909 | -2.000224238 | 0.114563708 | #N/A | #N/A |
| RNAseq.g18123.t1 | MFI2 | ABC transporter, phosphonate, periplasmic substrate-binding protein | 0.5443209609 | 0.04989535616 | 0.7900964494 | 0.08397191935 | 0.485248832 | 0.155344043 | 0.755081405 | 0.03525361 |
| RNAseq.g18955.t1 | SLC2A4 | hexose transmembrane transporter activity | 0.2245327474 | 0.7012837486 | 1.000591304 | 0.0473587749 | 0.129516894 | 0.883593125 | 0.799437416 | 0.157865358 |
| RNAseq.g19120.t1 | SLC7A1 | cationic amino acid transporter | -0.5808741534 | 0.6841130496 | -0.7893726297 | 0.6678814323 | -0.945658171 | 0.532502801 | #N/A | #N/A |
| RNAseq.g21401.t1 | ALDH1L2 | belongs to the aldehyde dehydrogenase family | -0.3433605877 | 0.3564437871 | 0.6121290045 | 0.207113614 | -0.227243893 | 0.655448822 | 0.571487108 | 0.177933785 |
| RNAseq.g22092.t1 | PLA2G4A | Phospholipase A2 | 5.438813521 | 1.49099225285309e-06 | #N/A | #N/A | 3.949441417 | 0.002439569 | #N/A | #N/A |
| RNAseq.g2272.t1 | amt-1 | ammonium transmembrane transporter activity | -1.423395665 | 0.2421831934 | 0.1247053985 | 0.9567840296 | -1.55035824 | 0.210769416 | 0.573488079 | 0.810436711 |
| RNAseq.g2273.t1 | amt-1 | ammonium transmembrane transporter activity | 1.583472166 | 0.2831512307 | -1.287094778 | 0.3980683981 | 1.139089227 | 0.40708172 | -1.753794834 | 0.101924799 |
| RNAseq.g23850.t1 | GAL3ST1 | galactose-3-O-sulfotransferase 1 | #N/A | #N/A | -1.695098228 | 0.07253930185 | -4.733769663 | 0.004915827 | #N/A | #N/A |
| RNAseq.g24789.t1 | HHIPL1 | scavenger receptor activity | 1.325626309 | 2.34247676473326e-06 | 0.5262107292 | 0.4957595876 | 1.290762827 | 8.28E-05 | 0.607872203 | 0.279633526 |
| RNAseq.g25818.t1 | H6PD | glucose dehydrogenase activity | 1.234212722 | 0.0600810626 | 0.07368833563 | 0.9814501808 | 1.261804696 | 0.110887111 | 0.082346852 | 0.979226836 |
| RNAseq.g25983.t1 | ALDH3B1 | Belongs to the aldehyde dehydrogenase family | 0.2779345276 | 0.3785917253 | -0.1474146286 | 0.9010443876 | 0.243183273 | 0.50969179 | -0.038299181 | 0.977631266 |
| RNAseq.g26193.t1 | ALOX5 | arachidonate 5-lipoxygenase activity | 1.11186483 | 0.00834980234 | 0.2339097903 | 0.9063667355 | 1.103266542 | 0.042275021 | 0.2343185 | 0.894018937 |
| RNAseq.g26193.t1 | ALOX5 | arachidonate 5-lipoxygenase activity | 1.11186483 | 0.00834980234 | 0.2339097903 | 0.9063667355 | 1.103266542 | 0.042275021 | 0.2343185 | 0.894018937 |
| RNAseq.g26356.t1 | PLA2G4A | Phospholipase A2 | 2.602717033 | 1.55303284894131e-31 | 1.438189377 | 0.0001201005651 | 2.717095174 | 3.55E-26 | 1.57697947 | 3.97E-09 |
| RNAseq.g26759.t1 | Nidogen-like | Nidogen-like | 5.375842255 | 1.78236625622357e-07 | #N/A | #N/A | 4.153739455 | 0.001787791 | #N/A | #N/A |
| RNAseq.g847.t1 | alh-8 | malonate-semialdehyde dehydrogenase (acetylating) activity | 0.1184058596 | 0.8392970046 | 0.699872256 | 0.1745603969 | 0.133542344 | 0.848595665 | 0.679171534 | 0.221952765 |
| RNAseq.g850.t1 | SLC29A1 | nucleoside transmembrane transporter activity | 0.1769067564 | 0.7586836549 | 1.171431267 | 0.003243844162 | -0.000254161 | 0.999538276 | 1.105936564 | 0.008136767 |
| TS.g1376.t2 | swt-4 | Sugar transporter | 0.944768975 | 0.01160404938 | 0.005874635552 | 0.9971933397 | 1.000755181 | 0.022270164 | -0.023048382 | 0.989988033 |
| TS.g1941.t1 | Nidogen-like | Nidogen-like | -0.9957716041 | 0.554862136 | -0.713186517 | 0.6395862741 | -1.321767424 | 0.41175008 | #N/A | #N/A |
| TS.g21811.t1 | Tnfrsf22 | TNFR/NGFR cysteine-rich region | 0.9589446835 | 1.87628801203033e-05 | 1.100350829 | 0.0001143400792 | 0.974994982 | 4.32E-05 | 0.978105368 | 0.000211621 |
| TS.g2203.t1 | SULT6B1 | Belongs to the sulfotransferase 1 family | -0.1652871635 | 0.8634833828 | -0.3959017982 | 0.8156384048 | -0.27749806 | 0.761502843 | -0.472959883 | 0.724330752 |
| TS.g27687.t1 | CHST13 | Sulfotransferase family | -1.371254788 | 0.1251855384 | -0.5706099238 | 0.8109495252 | -1.434466187 | 0.151973392 | #N/A | #N/A |
| TS.g7018.t1 | Cuzd1 | scavenger receptor activity | -1.264625755 | 0.03169019614 | -0.3357656514 | 0.7056352942 | -1.024710672 | 0.082929184 | -0.036048516 | 0.985876162 |
| TS.g7997.t1 | ALOX5 | arachidonate 5-lipoxygenase activity | 0.5525549355 | 0.2739983337 | 0.1957439471 | 0.9237595353 | 0.550902601 | 0.37526484 | 0.180435321 | 0.913596645 |
| TS.g8426.t1 | SLC6A19 | Transporter | -1.811642972 | 3.02697459111298e-05 | -0.4326550412 | 0.6620274656 | -1.725519568 | 4.67E-05 | -0.494421898 | 0.546096401 |

**Table S6.**

**Table S6.** *Pocillopora acuta* host RNA sequencing information including sample name, host genotype, raw read amount, reads mapping to the reference host genome, and percentage of cleaned reads mapped to the host genome.

| Sample_name | Genotype | Raw reads | Mapped reads | % Reads mapped |
|---|---|---|---|---|
| ECAH1 | 1:C | 11,651,308 | 6,598,310 | 56.63149579 |
| ECAH2 | 1:C | 8,220,385 | 4,627,006 | 56.28697439 |
| ECAH3 | 1:C | 10,103,799 | 5,455,834 | 53.99784774 |
| ECAH4 | 1:C | 10,899,798 | 4,690,582 | 43.03365989 |
| ECAH5 | 1:C | 6,604,874 | 3,125,814 | 47.32586874 |
| ECAH6 | 1:C | 8,089,163 | 3,858,819 | 47.7035634 |
| ECG1 | 2:D | 9,468,653 | 5,910,973 | 62.42675701 |
| ECG2 | 2:D | 8,967,591 | 5,923,270 | 66.05196423 |
| ECG3 | 2:D | 11,773,510 | 7,579,521 | 64.37775141 |
| ECG4 | 2:D | 6,659,588 | 4,023,941 | 60.42327243 |
| ECG5 | 2:D | 6,923,973 | 4,725,066 | 68.24212053 |
| ECG6 | 2:D | 6,852,396 | 3,528,913 | 51.49896474 |
| ORAG1 | 3:D | 6,698,441 | 4,850,973 | 72.41943312 |
| ORAPG1 | 3:D | 7,110,346 | 4,825,501 | 67.86590976 |
| ORAPG2 | 3:D | 7,984,566 | 5,249,451 | 65.74497599 |
| ORAPG3 | 3:D | 8,061,969 | 5,640,587 | 69.96537694 |
| ORAPG4 | 3:D | 8,804,806 | 4,752,895 | 53.98068964 |
| ORAPG5 | 3:D | 8,932,953 | 4,614,512 | 51.65718436 |
| ORAPG6 | 3:D | 6,383,454 | 3,344,739 | 52.39700952 |

**Table S7**

**Table S7.** *Pocillopora acuta* 16S rRNA sequencing information including sample name, host genotype, input reads, filtered reads, denoised reads, merged reads, tabled and nonchimeric reads.

| Sample name | Genotype | Sample name | Input reads | Filtered reads | Denoised reads | Merged reads | Tabled reads | Nonchimeric reads |
|---|---|---|---|---|---|---|---|---|
| ECAH1 | 1:C1 | ECAH1 | 5981 | 1848 | 1797 | 1792 | 1759 | 1759 |
| ECAH2 | 1:C2 | ECAH2 | 7283 | 3117 | 3104 | 3077 | 3077 | 3077 |
| ECAH3 | 1:C3 | ECAH3 | 3599 | 1219 | 1195 | 1194 | 1194 | 1194 |
| ECAH4 | 1:C4 | ECAH4 | 5586 | 2463 | 2448 | 2431 | 2413 | 2413 |
| ECAH5 | 1:C5 | ECAH5 | 10465 | 5309 | 5258 | 5181 | 5130 | 5130 |
| ECAH6 | 1:C6 | ECAH6 | 3687 | 1230 | 1216 | 1185 | 1185 | 1185 |
| ECG1 | 2:D1 | ECG1 | 9078 | 2854 | 2838 | 2795 | 2795 | 2741 |
| ECG2 | 2:D2 | ECG2 | 10986 | 5632 | 5590 | 5586 | 5586 | 5586 |
| ECG3 | 2:D3 | ECG3 | 5494 | 2316 | 2273 | 2251 | 2251 | 2251 |
| ECG4 | 2:D4 | ECG4 | 13199 | 8640 | 8624 | 8587 | 8582 | 8582 |
| ECG5 | 2:D5 | ECG5 | 6524 | 3367 | 3321 | 3321 | 3321 | 3321 |
| ECG6 | 2:D6 | ECG6 | 9674 | 6149 | 6130 | 6125 | 6094 | 6094 |
| ORAPG1 | 3:D1 | ORAPG1 | 4046 | 728 | 715 | 683 | 683 | 683 |
| ORAPG2 | 3:D2 | ORAPG2 | 5344 | 1608 | 1592 | 1546 | 1546 | 1546 |
| ORAPG3 | 3:D3 | ORAPG3 | 5211 | 1196 | 1172 | 1168 | 1138 | 1138 |
| ORAPG4 | 3:D4 | ORAPG4 | 7407 | 2494 | 2460 | 2432 | 2398 | 2398 |
| ORAPG5 | 3:D5 | ORAPG5 | 5277 | 1109 | 1088 | 1042 | 1042 | 1042 |
| ORAPG6 | 3:D6 | ORAPG6 | 5290 | 2990 | 2954 | 2933 | 2933 | 2933 |
